# Multi-scale electrophysiological characterization of human cardioids reveals progressive maturation

**DOI:** 10.64898/2026.09.29.755336

**Authors:** Varjany K. Vashanthakumar, Muhammad Naveed, Ekapaksi Wisnumurti, Nicolò Alerni, Lisa Conrad, Virginie Tissières, Johannes Tüchler, Iros Barozzi, Balázs Ördog, András Horváth, Marco Osterwalder, Katja E. Odening

## Abstract

**Background:** Human induced pluripotent stem cell-derived cardiomyocytes are valuable for cardiac disease modelling and drug testing but remain limited by immature structural and electrophysiological properties in conventional two-dimensional cultures. Three-dimensional cardiac models, including cardioids, provide a more physiological environment through multicellular organization and enable tissue-level functional analysis. Their electrophysiological properties and maturation, however, remain incompletely characterized.

**Methods:** We performed an integrated electrophysiological characterization of human cardioids over the first three weeks of maturation using sharp microelectrode recordings, calcium transient measurements, whole-cell patch clamp recordings, immunofluorescence, transcriptomic analysis, and pharmacological interventions.

**Results:** During maturation, cardioids exhibited progressive electrophysiological maturation, characterized by more physiological resting membrane potentials (−54.0±3.1mV to −68.1±1.2mV), increased action potential amplitude (73.5±4.8mV to 101.7±2.1mV) and upstroke velocity (44.3±5mV/ms to 86.6±8.4mV/ms), and shorter action potential duration (178.9±8.7ms to 144.0±4.8ms at 1Hz stimulation). These changes were associated with progressively increased I_Na_ density and enhanced *SCN5A* mRNA and Na_v_1.5 protein expression, consistent with progressive maturation of cardiac excitability. I_Kr_ current and *KCNH2*/hERG expression were already present at early maturation stages and remained stable, as evidenced by preserved sensitivity to dofetilide-blockade. Pharmacological inhibition of barium- and HMR-sensitive currents altered resting membrane and/or action potential characteristics in intact cardioids, whereas in isolated cardiomyocytes classical I_K1_ remained undetectable and I_Ks_ was small, highlighting a certain degree of persisting immaturity of the model.

**Conclusions:** These findings demonstrate that human cardioids undergo progressive electrophysiological maturation and provide complementary tissue- and cellular-level insights into cardiac function, establishing their suitability for investigating I_Na_ and I_Kr_-related cardiac pathologies.

## Introduction

Human induced pluripotent stem cell-derived cardiomyocytes (hiPSC-CMs) have transformed cardiac research by providing a renewable, patient-specific, and ethically accessible source of human cardiomyocytes for disease modelling, drug discovery, and personalized medicine (Takahashi and Yamanaka, 2006). Early studies demonstrated that patient-specific hiPSC-CMs successfully reproduced the electrophysiological phenotype of long QT syndrome *in vitro* (Moretti et al., 2010; Itzhaki et al., 2011). Since then, hiPSC-CMs have been widely used to study disease mechanisms, evaluate pro-arrhythmic drug effects, and support personalized therapeutic approaches (Joshi et al., 2024).

Despite these advances, however, conventional two-dimensional (2D) hiPSC-CM monolayers remain limited by immature structural and electrophysiological phenotype (Feyen et al., 2020; Odening et al., 2021). To better recapitulate native cardiac tissue, increasing attention has shifted toward three-dimensional (3D) cardiac models, including engineered heart tissues (Schaaf et al., 2014; Feyen et al., 2020), cardiac spheroids (Beauchamp et al., 2015), and cardioids (Ma et al., 2015; Hofbauer et al., 2021; Schmidt et al., 2023). Among these models, cardioids are self-organizing chamber-like tissues generated from hiPSCs with minimal external manipulation (Hofbauer et al., 2021). They recapitulate key features of early human cardiac development, including cavity formation, myocardial organization, spontaneous contraction, and myocyte-non-myocyte interactions. Moreover, basic left ventricle-type cardioids consisting of around 90 % cardiomyocytes can be generated reproducibly at scale within approximately one week (Hofbauer et al., 2021; Schmidt et al., 2023). Although the structural and developmental properties of cardioids are well established (Hofbauer et al., 2021, Schmidt et al., 2023), their electrophysiological characterization remains limited. Previous studies have largely focused on morphogenesis, cellular composition, and transcriptomics, with functional electrophysiological assessments being restricted to multielectrode arrays, optical calcium imaging, contraction analysis, and action potential recordings in isolated cardioid-derived cardiomyocytes after extended culture at one timepoint (Hofbauer et al., 2021; Schmidt et al., 2023). Consequently, tissue-level electrophysiology, pharmacological responses, cardiac ion currents, and electrophysiological maturation throughout cardioid development remain insufficiently understood. Furthermore, while hiPSC-CMs exhibit an immature electrophysiological phenotype characterized by depolarized resting membrane potentials, spontaneous automaticity, and immature cardiac ion currents (Veerman et al., 2015; Odening et al., 2021), it remains unclear whether the 3D architecture of cardioids enhances electrophysiological maturation.

In this study, we performed a comprehensive electrophysiological characterization of human cardioids over three weeks of maturation using sharp microelectrode recordings, calcium imaging and whole-cell patch clamp recordings, supported by immunofluorescence analysis and transcriptomic profiling. We assessed tissue-level electrical activity, pharmacological responses to ion channel blockers, and the maturation of major cardiac ion currents in line with ion channel expression dynamics, providing an integrated evaluation of electrophysiological maturation in human cardioids.

## Methods

### Generation and culture of left ventricular cardioids

Left ventricular (LV)-type cardioids were produced from the human induced pluripotent stem-cell (hiPSC) line AICS-0037-172:WTC-mEGFP-TNNI1-cl172, carrying a monoallelic mEGFP tag at the *TNNI1* locus, and following a protocol adapted from Hofbauer et al. (2021) with minor modifications (**Fig. 1**). The cell line was obtained from the Allen Institute for Cell Science Human iPSC Collection under a Uniform Biological Material Transfer Agreement with the Coriell Institute for Medical Research.

**Figure 1.**
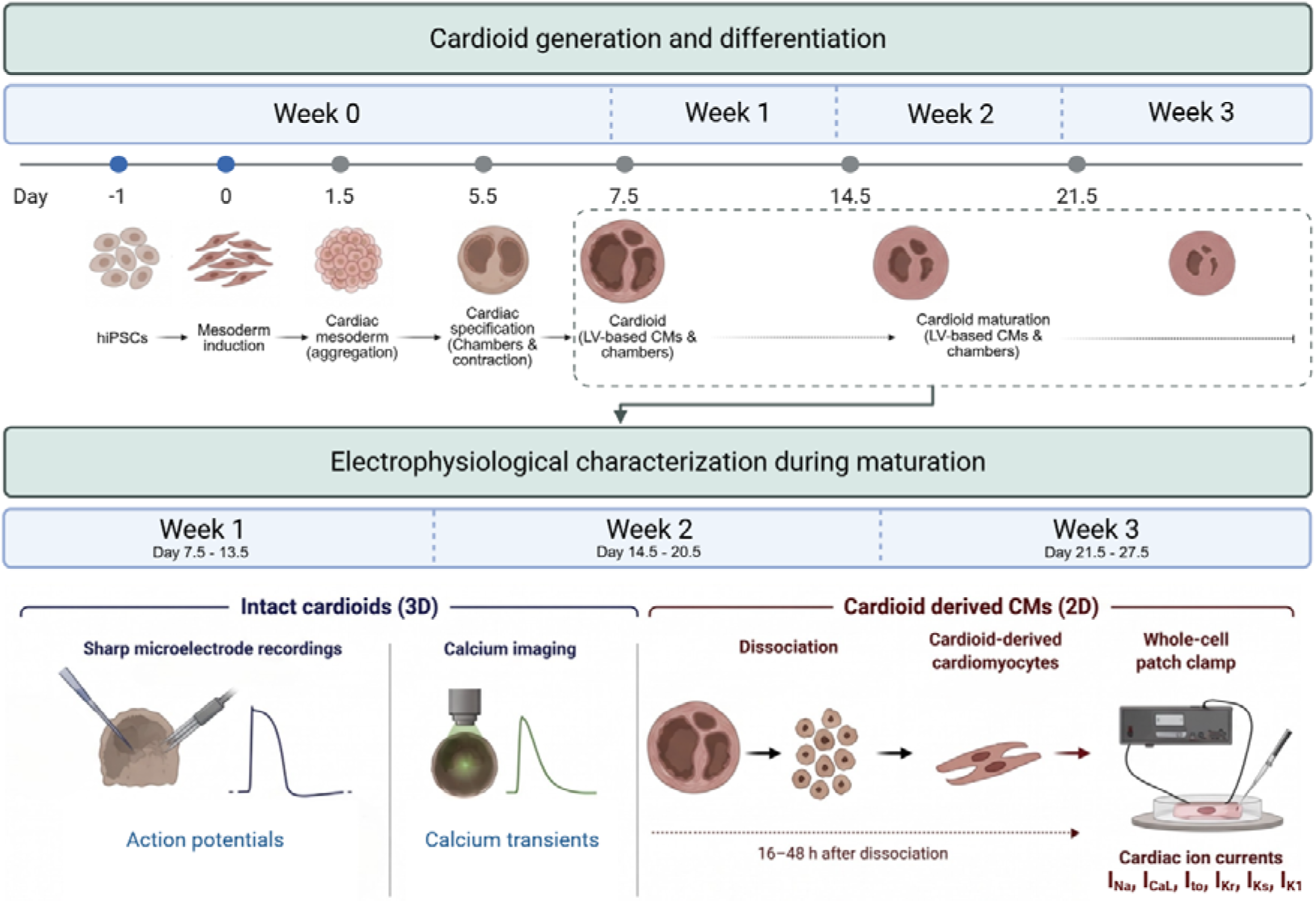
Experimental timeline illustrating cardioid generation and electrophysiological characterization. hiPSCs were differentiated toward the cardiac lineage by mesoderm induction for 36 h, followed by aggregation in U-shaped 96-well plates on day 1.5 to generate cardioids (Hofbauer et al., 2021). After sequential medium changes, cardioids were maintained in maintenance medium containing insulin from day 7.5 onwards. Electrophysiological experiments were performed during week 1 (days 7.5-13.5), week 2 (days 14.5-20.5), and week 3 (days 21.5-27.5). Action potentials and calcium transients were recorded from intact cardioids paced at 1 Hz using sharp micro-electrodes and calcium imaging. Cardioids were subsequently dissociated using the in-house developed LATLe enzyme mixture (Liberase, Accutase and TrypLE), and whole-cell patch clamp recordings were performed in isolated cardiomyocytes 16-48 h after dissociation at 37°C using Tyrode solution as the external solution.

### hiPSC maintenance

hiPSCs were maintained under feeder-free conditions on Matrigel-coated (Corning hESC-qualified Matrix, Merck, CLS354277) tissue culture plates in mTeSR™ Plus medium (StemCell Technologies, #100-0276). Cells were passaged every 3-4 days at approximately 80 % confluency using TrypLE™ Express (Gibco™, Fisher Scientific, #10043382). Following each passage, cultures were supplemented with 5-10 μM ROCK inhibitor Y-27632 (Bio-Techne, #1254/10) for less than 24 hours to enhance single-cell survival. hiPSCs were routinely tested and found negative for mycoplasma contamination.

### Two-dimensional mesoderm induction

For differentiation, hiPSCs were dissociated into single cells and seeded onto Matrigel-coated 24-well plates at 30,000 cells per well in mTeSR Plus medium supplemented with 10 μM ROCK inhibitor (Y-27632). Only cultures with greater than 90 % viability and homogeneous cell distribution were used for differentiation.

All differentiation media were based on chemically defined medium (CDM), consisting of a 1:1 mixture of IMDM (Gibco, Fisher Scientific, 11510596) and Ham’s F-12 GlutaMAX (Gibco, Fisher Scientific, 11514446), supplemented with 5 mg/ml bovine serum albumin (Bovogen, BSAS-AU), 1 μl/ml chemically defined lipid concentrate (Gibco, Fisher Scientific, 11548846), 15 μg/ml transferrin (Roche, Merck, 10652202001), and 0.04 μl/ml 1-thioglycerol (Merck, M6145-25ML).

Thirty hours after seeding, the medium was replaced with LV induction medium comprising CDM supplemented with 10 ng/ml BMP4, Bone morphogenetic protein 4 (R&D Systems, Bio-Techne, 314-BP-050), 5.5 ng/ml Activin A (Qkine, QK001-0025), 5.5 ng/ml FGF2 (R&D Systems, Bio-Techne, 3718-FB), 5 μM LY294002 (Tocris, Bio-Techne, 1130/5), and 3 μM CHIR99021 (Tocris, Bio-Techne, 4423/10). The start of LV induction was designated differentiation day 0 (D0). Cultures were maintained in LV induction medium for 36-40 h, until they formed a confluent mesodermal monolayer, typically exceeding 90 % confluency.

### Three-dimensional cardioid formation and maturation

At D1.5, mesodermal monolayers were dissociated with TrypLE Express for approximately 1 min and gently resuspended in LV CardMeso medium. LV CardMeso medium consisted of CDM supplemented with 10 μg/ml insulin (Roche, Merck, 11576497001), 10 ng/ml BMP4, 1.46 ng/ml FGF2, 2 μM Wnt-C59 (Tocris, Bio-Techne, 5148/10), 0.05 μM retinoic acid (Merck, R2625), and 5 μM ROCK inhibitor (Y-27632).

Cell suspensions were counted using Trypan Blue exclusion, and only preparations with greater than 90 % viability were used. Cells were seeded at 15,000 cells per well in ultra-low-attachment 96-well plates to induce three-dimensional aggregation. Plates were centrifuged at 140 g for 3 min to promote uniform spheroid formation.

Culture medium was exchanged daily by replacing 180 μl per well. On D2.5, the medium was changed to LV CardMeso medium without ROCK inhibitor, and cardioids were maintained under these conditions until D5.5. At D5.5, cardioids were transferred to LV CardMyo medium consisting of CDM supplemented with 10 μg ml−1 insulin, 10 μg/ml BMP4, and 1.46 ng/ml FGF2. LV CardMyo medium was refreshed daily until D7.5. Thereafter, cardioids were maintained in CDM supplemented with 10 μg/ml insulin, with medium changes every other day, until further experimentation (**Fig. 1**). A cardiac differentiation was verified by assessing expression profiles of the cardiomyocyte marker genes *MYH6*, *TNNT2*, *TNNI1* and *ACTN2* in hiPSCs (D0), mesodermal progenitors (D1.5), and cardioids at D5.5 and D7.5 (**Suppl. Fig. 1**).

### Electrophysiological evaluation of cardioids

Electrophysiological assessment was performed at both the tissue and cellular levels. In intact cardioids, conventional sharp microelectrode technique was used to measure action potentials (APs), while calcium transients were recorded optically during 1 Hz electrical pacing. At the cellular level in cardioid-derived cardiomyocytes, the whole-cell configuration of the voltage-clamp technique was used to characterize the major cardiac ion currents, including the sodium current (I_Na_), transient outward potassium current (I_to_), L-type calcium current (I_CaL_), rapid delayed rectifier potassium current (I_Kr_), slow delayed rectifier potassium current (I_Ks_), and inward rectifier potassium current (I_K1_).

Unless otherwise stated, all recordings were performed in HEPES-buffered Tyrode’s extracellular solution containing (mM): 135 NaCl, 0.4 NaH2PO4, 5 KCl, 10 HEPES, 1 MgCl2, 1.8 CaCl2, 10 glucose, 10 creatine, and 20 taurine (pH 7.4 adjusted with NaOH), as previously described (Nimani et al., 2024; Louradour et al., 2025). The use of a common extracellular solution across sharp microelectrode, calcium imaging, and patch clamp experiments facilitated direct comparison between experimental approaches.

### Conventional sharp microelectrode action potential recordings in intact cardioids

Prior to electrophysiological recordings, cardioids were transferred onto Matrigel-coated glass coverslips (15 mm diameter, Carl Roth, #P232.1) positioned within cloning cylinders (Pyrex®, Merck, #CLS31668). Cardioids were allowed to attach for 48-72 hours while maintaining their three-dimensional architecture. Week 1 (D7.5-D13.5) cardioids typically attached within 48 hours, whereas week 2 (D14.5-D20.5) and week 3 (D21.5-D27.5) cardioids generally required approximately 72 hours.

Conventional sharp microelectrode technique (Jost et al., 2013b) in intact cardioids were performed using a modified patch clamp setup adapted for extracellular stimulation and intracellular microelectrode recordings. APs were recorded using an Axopatch-200B amplifier coupled to a Digidata 1550B data acquisition system (Molecular Devices, CA, USA). Cardioids were electrically stimulated at 1 Hz using concentric bipolar stimulation electrodes (MicroProbes, Science Products GmbH, Germany). Electrical stimulation was delivered using an external pulse stimulator (MyoPacer, IonOptix LLC, USA, or Stimulus Isolator, World Precision Instruments, USA), with stimulation protocols generated through pClamp 11.1 software (Molecular Devices). Stimulation pulses were delivered for 2-4 ms at an amplitude of 20-30 V.

Glass microelectrodes were filled with 3 M KCl and had tip resistances of 15-30 MΩ. Recordings were performed in Tyrode’s solution at 37°C. To reduce contraction-induced motion artefacts, approximately half of the experiments were performed on cardioids incubated with 10 μM blebbistatin (Biozol, #B592490) for at least 40 minutes before recordings. As blebbistatin treatment did not significantly affect AP parameters (**Suppl. Fig. 2**), data obtained in the presence and absence of blebbistatin were pooled for subsequent analyses.

After stable baseline recordings had been obtained, cardioids were superfused with Tyrode’s solution containing the respective pharmacological inhibitor for 5-20 min, and electrophysiological responses to drug application (dofetilide (1 μM), HMR-1556 (1 μM), or Barium chloride (BaCl₂) (100 μM)) were recorded. If microelectrode impalement was lost during the recording, re-impalement was attempted. Recordings were accepted only when action potential characteristics after re-impalement differed by less than 5% from the preceding recording; otherwise, the experiment was excluded from further analysis.

Resting membrane potentials (RMP), action potential amplitude (APA), maximal upstroke velocity (dV/dt_max_) and action potential duration at 80% of repolarization (APD_80_) were analysed using custom-developed software, APEX (Action Potential Examination Platform; © 2025-2026, University of Bern, Institute of Physiology, Odening Lab; Vashanthakumar VK. & Odening KE., 2026; available at https://github.com/varjany-kunalini-vashanthakumar/APEX; DOI: 10.5281/zenodo.23026859).

### Calcium transient recordings in intact cardioids

Intracellular calcium transients were recorded from intact cardioids using the IonOptix fluorescence imaging system (IonOptix LLC, USA). Data acquisition and analysis were performed using IonWizard software (IonOptix LLC, USA) (as in Alerni et al., 2026; Louradour et al., 2025).

Prior to recordings, cardioids were attached to Matrigel-coated glass coverslips as described above and incubated for 15 minutes at room temperature with 1.5 μM Fura-2 AM (Abcam, UK).

Cardioids were electrically stimulated at 1 Hz using a MyoPacer field stimulator (IonOptix LLC, USA). Fura-2 was excited alternately at 340 nm and 380 nm, corresponding to Ca²⁺-bound and Ca²⁺-free dye, respectively, while emitted fluorescence was detected at 510 nm using a photomultiplier tube (PMT). Fluorescence signals were integrated using a Fluorescence System Interface (FSI).

Calcium handling parameters, including time-to-peak, maximal upstroke velocity (dcdt, ΔF/F0*ms^-1^), and calcium transient duration at 80 % recovery (CaD_80_), were analysed using IonWizard software.

### Cardioid dissociation

Cardioids were dissociated into single cardiomyocytes for whole-cell patch clamp recordings as follows. Briefly, cardioids were washed twice with phosphate-buffered saline (PBS) and incubated in PBS for 15 minutes to remove residual culture medium. For each experimental batch and maturation stage, cardioids were first mechanically fragmented and incubated at 37 °C in an enzyme mixture consisting of Liberase™ TL (∼2 WU, Merck, #5401020001), Accutase™ (StemCell Technologies, #07922), and TrypLE™ Express (2.5×, Gibco™, Fisher Scientific, #10043382) mixed at a volumetric ratio of 25:50:25, respectively. Approximately 500 μl of enzyme mixture was used per isolation, containing a varying number of cardioids at different maturation stages as cardioids became smaller and more compact with longer culture (20-25 cardioids at week 1, 30-35 at week 2, and 40-45 at week 3).

The digestion duration was adjusted according to maturation stage (15 minutes for week 1 cardioids, 20-25 minutes for week 2 cardioids, and 35-40 minutes for week 3 cardioids). Mechanical trituration was performed periodically using sterile plastic pipettes to facilitate dissociation. The remaining undigested tissue fragments were transferred into fresh enzyme solution and further incubated until complete dissociation was achieved.

The dissociation reaction was stopped by addition of maintenance medium supplemented with 5 μM ROCK inhibitor Y-27632 (Tocris, Bio-Techne, #1254/10). Cells were collected by centrifugation, resuspended in fresh ROCK inhibitor-containing maintenance medium, and counted. Approximately 75,000-100,000 cells were seeded onto 15 mm diameter Matrigel-coated glass coverslips depending on cell yield. Cells were maintained in medium containing ROCK inhibitor for 3 hours to promote attachment, after which the medium was replaced with maintenance medium without ROCK inhibitor. Whole-cell electrophysiological recordings were performed 16-48 hours after dissociation.

### Whole-cell Patch clamping

The membrane currents were recorded using either an Axopatch-200B amplifier coupled to a Digidata 1550B digitizer (Molecular Devices, USA) or a HEKA EPC10 Plus amplifier (HEKA/Multi Channel Systems, Germany) using the patch clamp technique in whole-cell configuration as described in (Lemoine et al., 2017; Horváth et al., 2018; Topal et al., 2021). Voltage protocols were generated and data were acquired using pClamp 11.1 (Molecular Devices) or PatchMaster software (HEKA), respectively.

Patch pipettes were fabricated from borosilicate glass capillaries using a DMZ Universal Electrode Puller (Munich, Germany) and had resistances of 1–3 MΩ after filling with internal solution. All recordings were performed at physiological temperature (37 °C). Current amplitudes were normalized to membrane capacitance and expressed as current density (pA/pF).

For recordings of I_Na_, I_to_, I_Kr_, I_Ks_, and I_K1_, the intracellular pipette solution contained (mM): 125 KCl, 5 NaCl, 1 MgCl2, 5 MgATP, 10 HEPES, and 5 EGTA (pH 7.2 adjusted with KOH).

For I_CaL_ recordings, the intracellular solution contained (mM): 130 CsCl, 10 TEA-Cl, 1 MgCl2, 5 MgATP, 2 EGTA, 1 CaCl2, 10 HEPES, and 0.3 GTP-Tris (pH 7.2 adjusted with CsOH).

To pharmacologically isolate individual currents, 1 μM nisoldipine was added to the HEPES-buffered Tyrode’s extracellular solution during recordings of I_Na_, I_to_, I_Kr_, I_Ks_, and I_K1_ to suppress I_CaL_. During I_Kr_ recordings, I_Ks_ was inhibited with 0.5 μM HMR-1556. For I_Ks_ recordings, I_Kr_ was inhibited using 0.1 μM dofetilide, and the bath solution was supplemented with 2 μM forskolin to enhance I_Ks_ activation. I_CaL_ recordings were performed in HEPES-buffered Tyrode’s solution supplemented with 3 mM 4-aminopyridine (4-AP) to suppress outward potassium currents.

I_Na_ was recorded from a holding potential of −110 mV using 30 ms depolarizing voltage steps from −80 to +60 mV in 5 mV increments, with an interpulse interval of 1 s. P/4 online leak subtraction was applied using four leak pulses, with leak pulses ranging from −92.5 to −136 mV, a 200-μs leak delay, a scaling factor of −0.25, and a leak holding potential of −100 mV.

I_CaL_ was recorded with cells held at −80 mV followed by a 100 ms prepulse to −40 mV to inactivate sodium channels. Cells were then subjected to 300 ms depolarizing voltage steps from −40 to +30 mV in 5-mV increments, delivered every 5 s.

I_K1_ was recorded from a holding potential of −90 mV using 300 ms voltage steps ranging from −110 to +60 mV in 10 mV increments at an interpulse interval of 3 seconds. A 50 ms prepulse to −40 mV was applied to eliminate sodium current contamination.

I_to_ was recorded from a holding potential of −90 mV using 300 ms depolarizing voltage steps from −50 to +30 mV in 10 mV increments at an interpulse interval of 3 seconds. A 50 ms prepulse to −40 mV was applied before each test pulse to inactivate sodium channels.

I_Kr_ was activated by 1-second-long voltage steps applied from a holding potential of −80 mV at an interpulse interval of 20 seconds between −50 to +30 mV in 10mV increments, following a 150 ms prepulse to −40 mV to inactivate sodium channels. After each voltage step, tail currents were observed at −40 mV (2 seconds).

I_Ks_ was recorded from a holding potential of −80 mV following a 50 ms prepulse to −40 mV. Currents were elicited using 5-second depolarizing voltage steps from −30 to +30 mV, and steady-state current amplitudes were measured at the end of each depolarizing pulse.

The identity of I_Na_, I_to_, I_Kr_, I_K1_ and I_CaL_ was further verified by pharmacological blocking experiments using mexiletine (100 μM), 4-AP (3 mM), dofetilide (1 μM), BaCl₂ (1 mM), and nisoldipine (1 μM), respectively, applied at the end of the recording.

### Immunofluorescence

Cardioids were harvested at day 7.5 (week 1), day 14.5 (week 2) and day 21.5 (week 3) for immunofluorescence analysis. Staining was performed as previously described, with minor modifications (Abassah-Oppong et al., 2024). Cardioids were fixed in 4 % paraformaldehyde (PFA; Merck, P6148) in PBS for 30 min at room temperature with gentle agitation, washed three times in PBS, and cryoprotected in 30 % (w/v) sucrose in PBS at 4□°C until fully infiltrated, as determined by sample sinking.

Cardioids were embedded in OCT compound (VWR, 361603E) in cryomolds, oriented individually, and rapidly frozen on a metal surface pre-cooled with liquid nitrogen. Embedded samples were stored at −80□°C until sectioning. Frozen cardioids were sectioned at 12□µm using a CM1950 cryostat (Leica Microsystems), and sections were mounted on Superfrost Plus slides and stored at −80□°C until use.

Before staining, cryosections were equilibrated to room temperature for 5-10□min and washed three times with PBS. Sections were permeabilized in 0.2 % Triton X-100 in PBS for 20□min, washed three times in PBS, and blocked for 1□h at room temperature in 1% bovine serum albumin (BSA) in PBS containing 0.1 % Tween-20 (PBT). Sections were then incubated overnight at 4□°C with primary antibodies diluted 1:250 in 1% BSA in PBS. Primary antibodies were directed against hERG/KCNH2 (rabbit; Alomone Labs, APC-062), Nav1.5/SCN5A (rabbit; Cell Signaling Technology, 14421T), Cav1.2/CACNA1C (rabbit; Alomone Labs, ACC-003) and GFP (chicken; Invitrogen, A10262) for detection of the TNNI1-GFP reporter.

Following primary antibody incubation, sections were washed three times for 10□min in PBS and once in PBT. Sections were incubated for 1□h at room temperature with species-appropriate fluorophore-conjugated secondary antibodies diluted 1:1,000 in 1 % BSA in PBS: goat anti-chicken Alexa Fluor 488 (Invitrogen, A11039) and donkey anti-rabbit Alexa Fluor 647 (Invitrogen, A31571). Sections were washed three times in PBS, counterstained with DAPI (1□µg/ml; Merck, D9542) for 10□min, washed three additional times for 10□min in PBS, and mounted in Mowiol 4-88 mounting medium (Merck, 475904). Images were acquired using an LSM 980 confocal microscope (Zeiss). Images were processed using ImageJ (v1.54f). Brightness and contrast adjustments were applied uniformly to all samples within an experiment using Adobe Photoshop.

### RNA sequencing

Three independent biological replicates were collected for each developmental stage per batch (batch 1: D0, D1.5, D7.5; batch 2: D7.5, D14.5, D21.5). For hiPSC, at least 750’000 cells were collected per replicate, and at least 500’000 cells were collected per replicate for the day 1.5 (D1.5) mesoderm prior to aggregation. For each cardioid replicates, 8 cardioids were pooled at day 7.5 (D7.5, week 1), day 14.5 (D14.5, week 2), and day 21.5 (D21.5, week 3). Cardioids were transferred to microcentrifuge tubes, briefly rinsed with PBS, flash-frozen in liquid nitrogen and stored at −80 °C until RNA extraction. Total RNA was isolated using the RNeasy Micro Kit (Qiagen, cat. no. 74004) according to the manufacturer’s instructions. Cardioids were lysed directly in the supplied lysis buffer and homogenized by repeated pipetting. RNA was eluted in 30 μl nuclease-free water and submitted to Macrogen Europe for analysis and sequencing. RNA concentration and integrity (RIN ≥ 9) were assessed using a 2200 TapeStation system and RNA ScreenTape (Agilent, cat. no. 5067-5576).

Strand-specific poly(A)-selected RNA-sequencing libraries were prepared from 1 μg total RNA using the TruSeq Stranded mRNA Library Preparation Kit (Illumina, cat. no. 20020595) and TruSeq RNA UD Indexes (Illumina, cat. no. 20022371), according to the manufacturer’s instructions. Library size distributions were assessed using a 2200 TapeStation system with D1000 ScreenTape (Agilent, cat. no. 5067-5582), and libraries were quantified by qPCR according to the Illumina qPCR Quantification Protocol Guide using a Roche LightCycler instrument. Equimolar pooled libraries were sequenced on an Illumina NovaSeq X Plus platform in paired-end mode (2 × 150 bp), generating approximately 20 million read pairs per sample.

### RNA-seq data processing and differential expression analysis

Raw paired-end reads were processed with the nf-core/rnaseq pipeline (version 3.14.0) (Ewels et al., 2020) implemented in Nextflow (Di Tommaso et al., 2017). Reads were quality-controlled and adapter-trimmed, aligned to the human reference genome (Ensembl GRCh38, primary assembly) with STAR (Dobin et al., 2013), and transcript-level abundances were quantified with Salmon (Patro et al., 2017) against the Ensembl GRCh38 release 112 annotations (**Suppl. Tables 3, 4**).

Transcript-level estimates were summarised to gene level in R using tximport (Soneson et al., 2015). Differential expression analysis was performed with DESeq2 (Love et al., 2014), including batch correction, modelling developmental stage as the sole variable of interest (D7.5, D14.5, D21.5) (**Suppl. Table 4, 5**). All pairwise contrasts between stages (D7.5, D14.5, D21.5) were tested by sequentially re-levelling the stage factor and refitting the model. Log2 fold-change estimates were shrunk using the apeglm estimator (Zhu et al., 2019). Genes with a Benjamini-Hochberg adjusted p-value < 0.05 and an absolute shrunken log2 fold change > 0.58 (1.5 linear fold) were considered differentially expressed. All analyses were conducted in R (version 4.3.1).

### Statistical analysis

Statistical analyses were performed using GraphPad Prism version 11.0.1 (GraphPad Software, USA). Data normality was assessed using the Shapiro-Wilk test. Normally distributed datasets were analysed using one-way analysis of variance (ANOVA) followed by Tukey’s or Dunnett’s multiple-comparisons test, as appropriate. Non-normally distributed data were analysed using the Kruskal-Wallis test with Dunn’s multiple-comparisons correction. Comparisons between paired baseline and drug-treated recordings were performed using paired Student’s *t*-tests or the Wilcoxon matched-pairs signed-rank test, as appropriate. Data are presented as mean ± SEM. All statistical tests were two-tailed, and differences were considered statistically significant at *p* < 0.05.

## Results

### Progressive maturation of AP properties in intact cardioids

To assess electrophysiological properties in human cardiac organoids, we generated left ventricular (LV)-type cardioids using an established protocol (Hofbauer et al., 2021) (**Figure 1**). At day 7.5 (D7.5), cardioids showed strongly elevated expression of the cardiomyocyte marker genes *MYH6, TNNT2, TNNI1 and ACTN2* as compared to D1.5 mesodermal progenitors and TNNI1-positive cells remained the predominant population throughout D7.5, D14.5 and D21.5 (**Suppl. Fig. 1**).

To determine whether LV-type cardioids undergo electrophysiological maturation during early development, we characterized APs during cardioid maturation at weeks 1-3 (**Figure 2**, **Suppl. Table 1**) using sharp microelectrode recordings.

**Figure 2.**
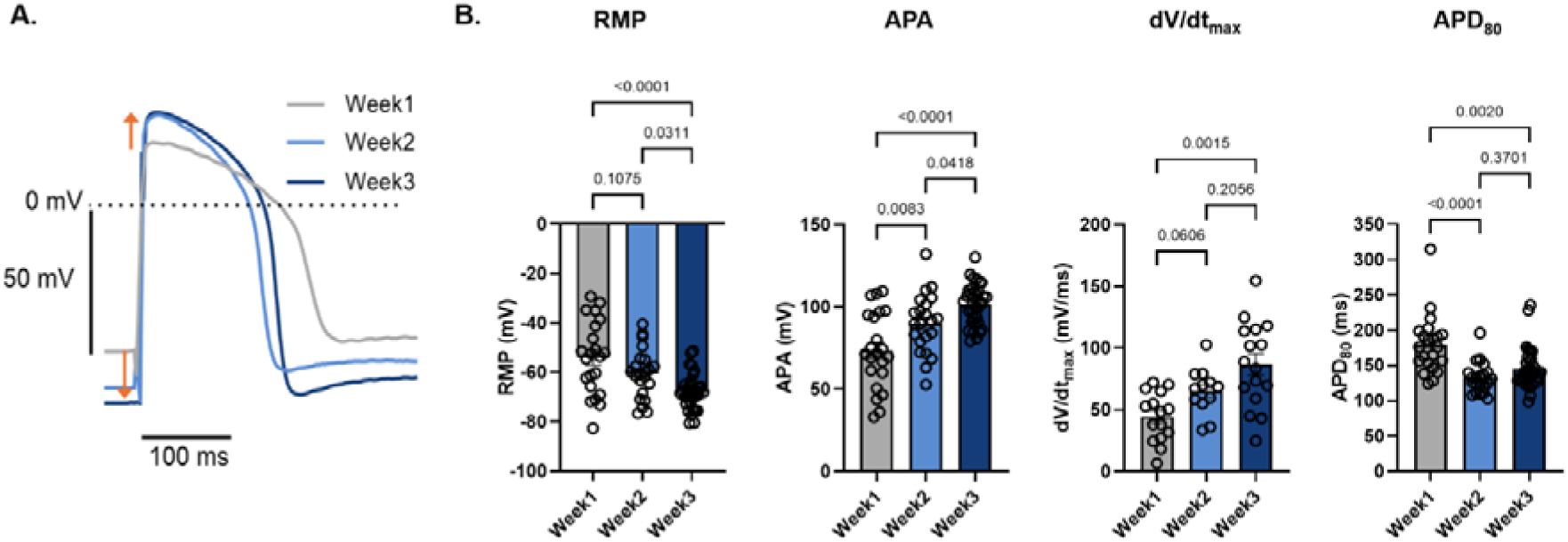
Progressive maturation of action potentials in intact cardioids. **A.** Representative action potentials recorded from week 1 (grey), week 2 (light blue) and week 3 (dark blue) cardioids. **B.** Bar graphs indicating statistical analyses of differences in resting RM), APA, dV/dt_max_ and APD_80_. RMP became progressively more negative (indicated by ↓), and APA and dV/dt_max_ increased significantly with maturation (indicated by ↑), whereas APD_80_ progressively shortened from week 1 to week 3. Data are presented as mean ± SEM. Statistical significance was defined as *p*<0.05. For RMP, APA and APD recordings n=22 cardioids were used at week 1, n=22 at week 2, and n=34 at week 3. Because the upstroke was frequently affected by pacing stimulation artefacts, only traces with a clearly identifiable upstroke were included in the dV/dt_max_ analysis, resulting in n=16, 12, and 15 cardioids at weeks 1, 2, and 3, respectively

RMP showed a progressive shift toward more negative potentials, from −54.0±3.1 mV at week 1 to −60.6±2.1 mV at week 2 and −68.1±1.2 mV at week 3 (**Figure 2B**), accompanied by increasing APA, from 73.5±4.8 mV at week 1 to 89.8±3.8 mV at week 2 and 101.7±2.1 mV at week 3 (**Figure 2B**) and increasing upstroke velocity dV/dt_max_ from 44.3±5.0 mV/ms at week 1 to 65.1±5.2 mV/ms at week 2 and 86.6±8.4 mV/ms at week 3 (**Figure 2B**). Because the upstroke was frequently affected by pacing stimulation artefacts, recordings were individually screened and only traces with a clearly identifiable upstroke were included in the dV/dt_max_ analysis, resulting in n = 16, 12, and 15 cardioids at weeks 1, 2, and 3, respectively development (**Suppl. Table 1B**). APD_80_ shortened gradually during maturation, decreasing from 178.9±8.7 ms at week 1 to 144.0±4.8 ms at week 3 (**Figure 2B**).

### Functional repolarizing currents contribute to APD in intact cardioids

The contribution of major cardiac repolarizing currents to APD in intact cardioids was assessed using pharmacological inhibitors during sharp microelectrode recordings (**Figure 3**, **Table 1**).

**Figure 3.**
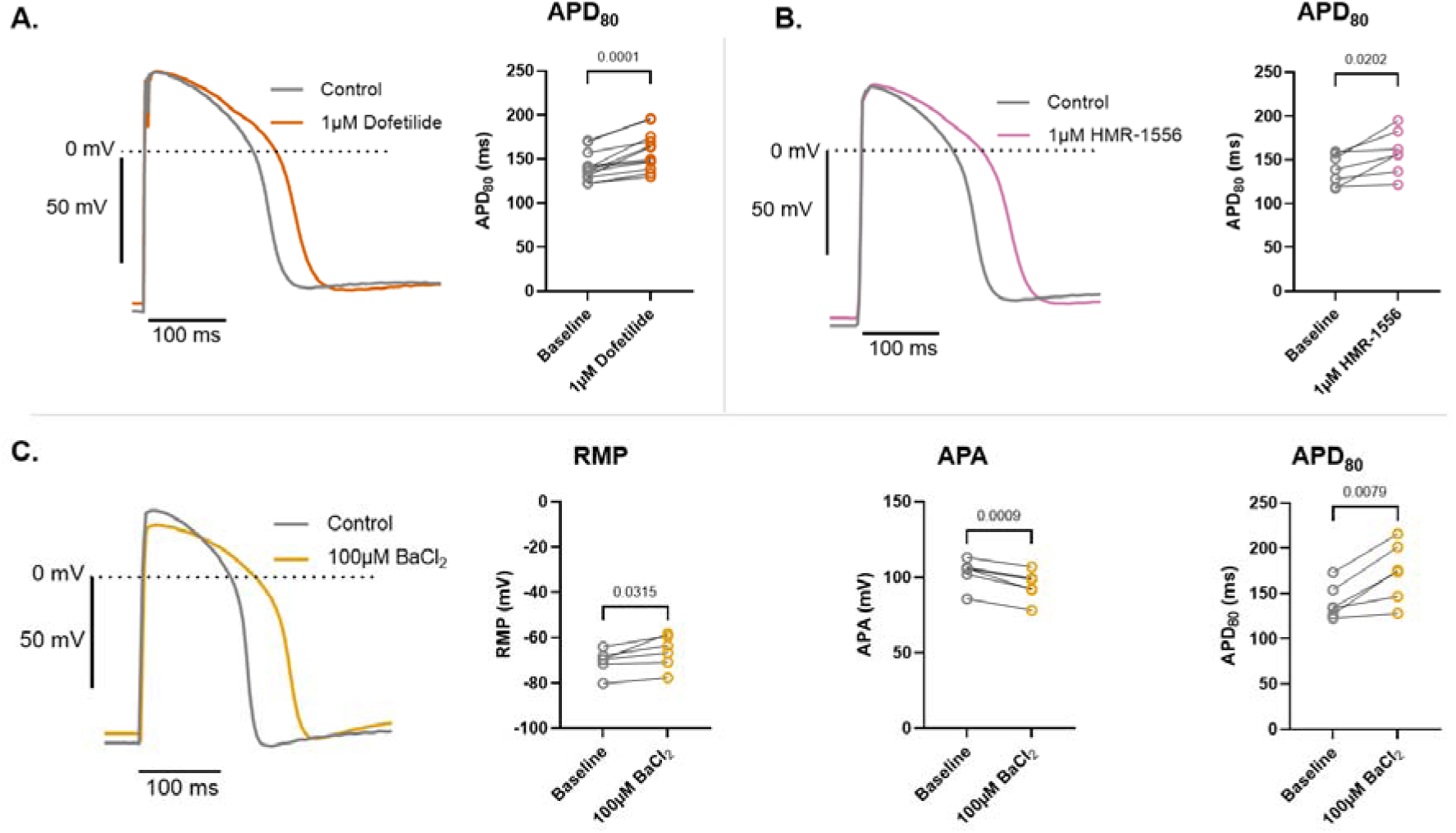
Pharmacological characterization of repolarization in intact cardioids. **A.** Effect of the selective I_Kr_ blocker dofetilide (1 μM) on APD_80_ in cardioids from week 2 (n=4), and week 3 (n=8). Dofetilide significantly prolonged APD_80_ throughout maturation. Individual cardioids are connected with line. **B.** Effect of the selective I_Ks_ blocker HMR-1556 (1 μM) on APD_80_ in cardioids in cardioids from week 2 (n=4), and week 3 (n=3). HMR-1556 significantly prolonged APD_80_, indicating functional contribution of I_Ks_ to repolarization. **C.** Effect of BaCl_2_ (100 μM) on action potential characteristics in week 2 (n = 1) and week 3 (n=5) cardioids. BaCl_2_ significantly depolarized the RMP, reduced APA, and prolonged APD_80_ in week 2 and week 3 cardioids, consistent with the contribution of one or more barium-sensitive currents to RMP and APD (i.e.I_K1_, I_K2P_). Data are presented as mean ± SEM. Paired t-test. Statistical significance was defined as p<0.05.

**Table 1.** Pharmacological effects of ion channel blockers on action potential characteristics in intact cardioids. RMP, APA, and APD_80_ were measured before (Control) and after application of dofetilide (1 μM), HMR-1556 (1 μM), or BaCl₂ (100 μM). N indicates the number of independent differentiation batches and n the number of cardioids. Data are presented as mean ± SEM. Statistical comparisons between control and drug-treated conditions were performed using a paired Student’s *t*-test. *\*p*<0.05, *\*\*p*<0.01, *\*\*\*p*<0.001 versus the corresponding control.

| Drug | Batch (N) | Cardioids (n) | RMP (mV) | APA (mV) | APD<sub>80</sub> (ms) |
| --- | --- | --- | --- | --- | --- |
| Control | N=3 | n=12 | -68.8±2.3 | 107.3±5.0 | 141.5±4.5 |
| 1 μM Dofetilide | N=3 | n=12 | -64.5±2.8 | 98.4±4.6 | 159.4±6.1*** |
| Drug | Batch (N) | Cardioids (n) | RMP (mV) | APA (mV) | APD<sub>80</sub> (ms) |
| Control | N=3 | n=7 | -67.2±3.4 | 95.8±5.0 | 145.6±7.1 |
| 1 μM HMR | N=3 | n=7 | -68.5±2.6 | 95.4±5.8 | 158.7±8.8* |
| Drug | Batch (N) | Cardioids (n) | RMP (mV) | APA (mV) | APD<sub>80</sub> (ms) |
| Control | N=2 | n=6 | -70.5±2.0 | 103.2±3.5 | 140.9±7.1 |
| 100 μM BaCl<sub>2</sub> | N=2 | n=6 | -66.0±2.7* | 94.5±3.0*** | 173.6±12.2** |

Perfusion with the I_Kr_ blocker dofetilide (1 μM) was applied to cardioids from week 2 (n=4), and week 3 (n=8). Dofetilide markedly prolonged APD_80_ without significantly affecting RMP or APA (**Figure 3A**, **Table 1**). Across all maturation stages, dofetilide prolonged APD_80_ from 141.5±4.5 ms at baseline to 159.4±6.1 ms (*p*=0.0001), confirming the presence of functional I_Kr_ in cardioids.

Similarly, inhibition of I_Ks_ with HMR-1556 (1 μM), performed in cardioids from week 2 (n=4), and week 3 (n=3), resulted in prolonged APD_80_ without significantly affecting RMP or APA (**Figure 3B**, **Table 1**). APD_80_ increased from 145.6±7.1 ms at baseline to 158.7±8.8 ms (*p*<0.05), demonstrating that I_Ks_ contributes to repolarization in intact cardioids. This contribution was further supported by an additional experiment in which sequential application of dofetilide followed by HMR-1556 in cardioids from week 2 (n=2) and week (n=3) resulted in an additional APD_80_ prolongation compared with dofetilide alone (*p*<0.05; **Suppl Fig. 2**).

To investigate the potential contribution of the inward rectifier potassium current I_K1_ or other barium-sensitive currents to the RMP, 100 μM BaCl₂ (**Figure 3C**, **Table 1**) was applied to cardioids from week 2 (n=1) and week 3 (n=5). BaCl₂ application significantly depolarized the RMP from −70.5±2.0 mV to −66.0±2.7 mV (*p*<0.05), reduced APA from 103.2±3.5 mV to 94.5±3.0 mV (*p*<0.001), and prolonged APD_80_ from 140.9±7.1 ms to 173.6±12.2 ms (*p*<0.01) (**Figure 3C**, **Table 1**). These findings indicate the presence of barium-sensitive currents that contribute to RMP regulation and AP repolarization in intact cardioids.

### Calcium handling improves during cardioid maturation

To investigate whether electrophysiological maturation was accompanied by improvements in calcium handling, calcium transient recordings were performed in intact cardioids at weeks 1-3 (**Figure 4, Suppl. Table 2**).

**Figure 4.**
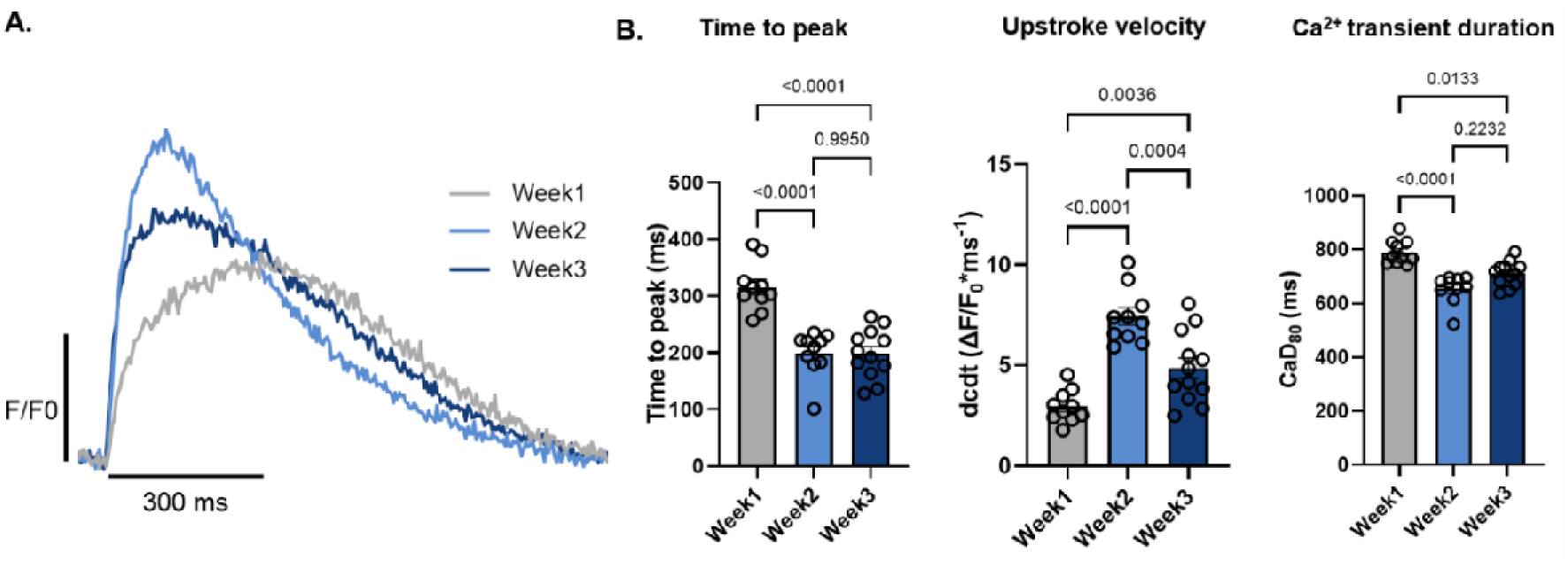
Calcium handling matures during cardioid development. **A.** Representative calcium transient recordings from week 1, week 2 and week 3 cardioids. **B.** Bar graphs indicating statistical analyses of differences in time-to-peak, upstroke velocity (dcdt) and calcium transient duration at 80% recovery (CaD_80_). Time-to-peak and CaD_80_ decreased with maturation, whereas upstroke velocity increased. n=10 cardioids per week. Data are presented as mean ± SEM. Statistical significance was defined as *p*<0.05.

Calcium transients demonstrated faster activation kinetics and shorter transient duration during maturation as indicated in the representative calcium transient traces (**Figure 4A**). Quantitative analysis revealed that time to peak significantly decreased from 315.8±12.8 ms at week 1 to 215.4±10.5 ms at week 3 (*p*<0.0001; **Figure 4B**), while dcdt increased from 3.0±0.2 ΔF/F0*ms^-1^ at week 1 to 4.9±0.6 ΔF/F0*ms^-1^ at week 3 (*p*<0.01; **Figure 4B**). Furthermore, CaD_80_ progressively shortened from 790.0±13.4 ms at week 1 to 720.9±14.5 ms at week 3 (*p*<0.05) (**Figure 4B**).

### Progressive increase in I_Na_ contributes to AP maturation, while I_Kr_ and I_CaL_ remain stable during cardioid development

To identify the ion currents underlying the AP maturation observed in intact cardioids, conventional whole-cell patch clamp recordings were performed in isolated cardioid-derived cardiomyocytes (**Figure 5**).

**Figure 5.**
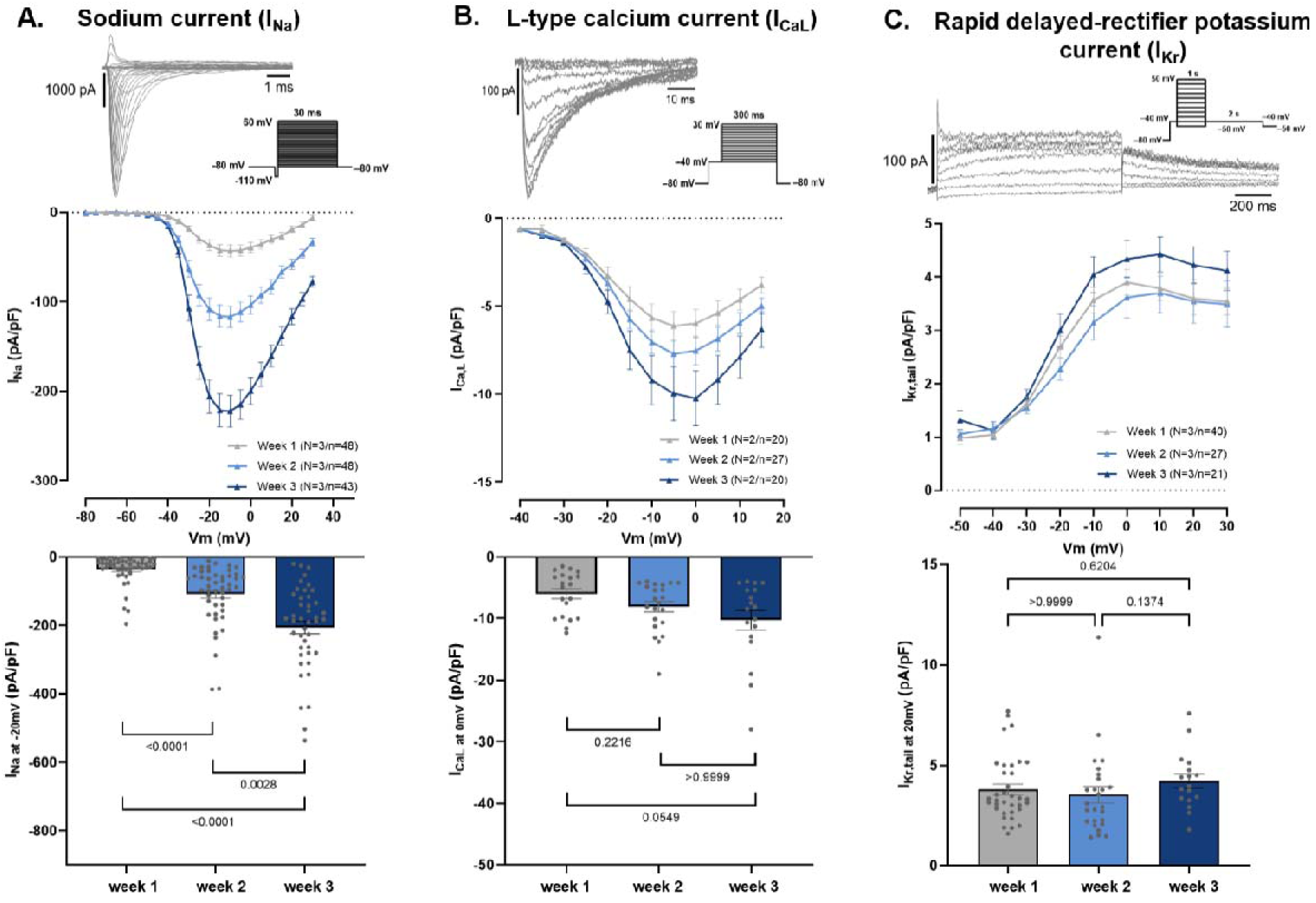
Changes of major cardiac ion currents during cardioid maturation. **A.** Sodium current (I_Na_). Representative current trace (top), current-voltage relationship (middle) and current density at −20 mV (bottom). Peak I_Na_ density increased significantly from week 1 to week 3. **B.** L-type calcium current (I_CaL_). Representative current trace (top), current-voltage relationship (middle) and current density at 0 mV (bottom). I_CaL_ was detected throughout maturation and showed a tendency to increase over time (p=0.0549). **C.** Rapid delayed rectifier potassium current (I_Kr_). Representative current trace (top), current-voltage relationship (middle) and current density at +30 mV (bottom). I_Kr_ was consistently detected and remained stable throughout maturation. Data are presented as mean ± SEM. N indicates the number of differentiation batches and n the number of isolated cardiomyocytes.

I_Na_ increased markedly throughout development (**Figure 5A**). Accordingly, peak current density (**Figure 5A**, lower panel) increased from −36.6±5.9 pA/pF at week 1 to −108.3±12.4 pA/pF at week 2 (*p*<0.0001) and −206.0±18.5 pA/pF at week 3 (*p*<0.0001), consistent with the progressive increase in APA and dV/dt_max_ observed in intact cardioids. In parallel, the proportion of cardiomyocytes exhibiting measurable I_Na_ increased during maturation (**Suppl. Fig, 4A**). Cells without detectable sodium current decreased from 21% at week 1 to 10% at week 2 and were absent by week 3 (**Suppl. Fig. 4A**). Conversely, the proportion of cells displaying very large sodium currents that precluded reliable voltage control increased from approximately 5% at week 1 to 27% at week 3 (**Suppl. Fig. 4A**).

I_to_ was detected throughout cardioid maturation (**Suppl. Fig. 5A**), exhibiting a non-significant upward trend over time.

I_CaL_ was detected at all developmental stages (**Figure 5B**). Peak current density measured at 0 mV (**Figure 5B**) showed a gradual increase from −6.0±0.8 pA/pF at week 1 to −8.1±0.8 pA/pF at week 2 and −10.2±1.5 pA/pF at week 3 (*p*=0.055).

I_Kr_ was consistently detected at all developmental stages (**Figure 5C**). Current density measured at +20 mV (**Figure 5C**) remained stable from 3.8±0.2 pA/pF at week 1 to 3.5±0.4 pA/pF at week 2 and 4.2±0.3 pA/pF at week 3, indicating that functional I_Kr_ is established early and maintained throughout cardioid maturation.

I_Ks_ steady current (**Suppl. Fig. 5B**) could only be detected in 58.5% (38 of 65) patched cardioid-derived cardiomyocytes (**Suppl. Fig. 4B**). Characteristic I_Ks_ tail currents could not be reliably detected in isolated cardioid-derived cardiomyocytes in our experimental conditions (**Suppl. Fig. 5B**).

Similarly, classical I_K1_ remained undetectable in isolated cardioid-derived cardiomyocytes (**Suppl. Fig. 4C**) across all maturation stages despite repeated recordings and the progressively more negative resting membrane potentials in intact cardioids (**Figure 2**).

### Increasing and stable ion-channel expression patterns accompany electrophysiological maturation in cardioids

To determine whether electrophysiological maturation was accompanied by changes in the expression of key ion channels, we performed immunofluorescence (IF) staining (**Figure 6A-C**) and bulk RNA sequencing (RNA-seq) in cardioids at D7.5, D14.5, and D21.5 (**Figure 6D, Suppl. Tables 3, 4**) and integrated the latter with the results from our previous RNA-seq experiment (D0, D1.5, D7.5) (see Methods) (**Suppl. Fig. 1A**, **Suppl. Tables 3, 4**).

**Figure 6.**
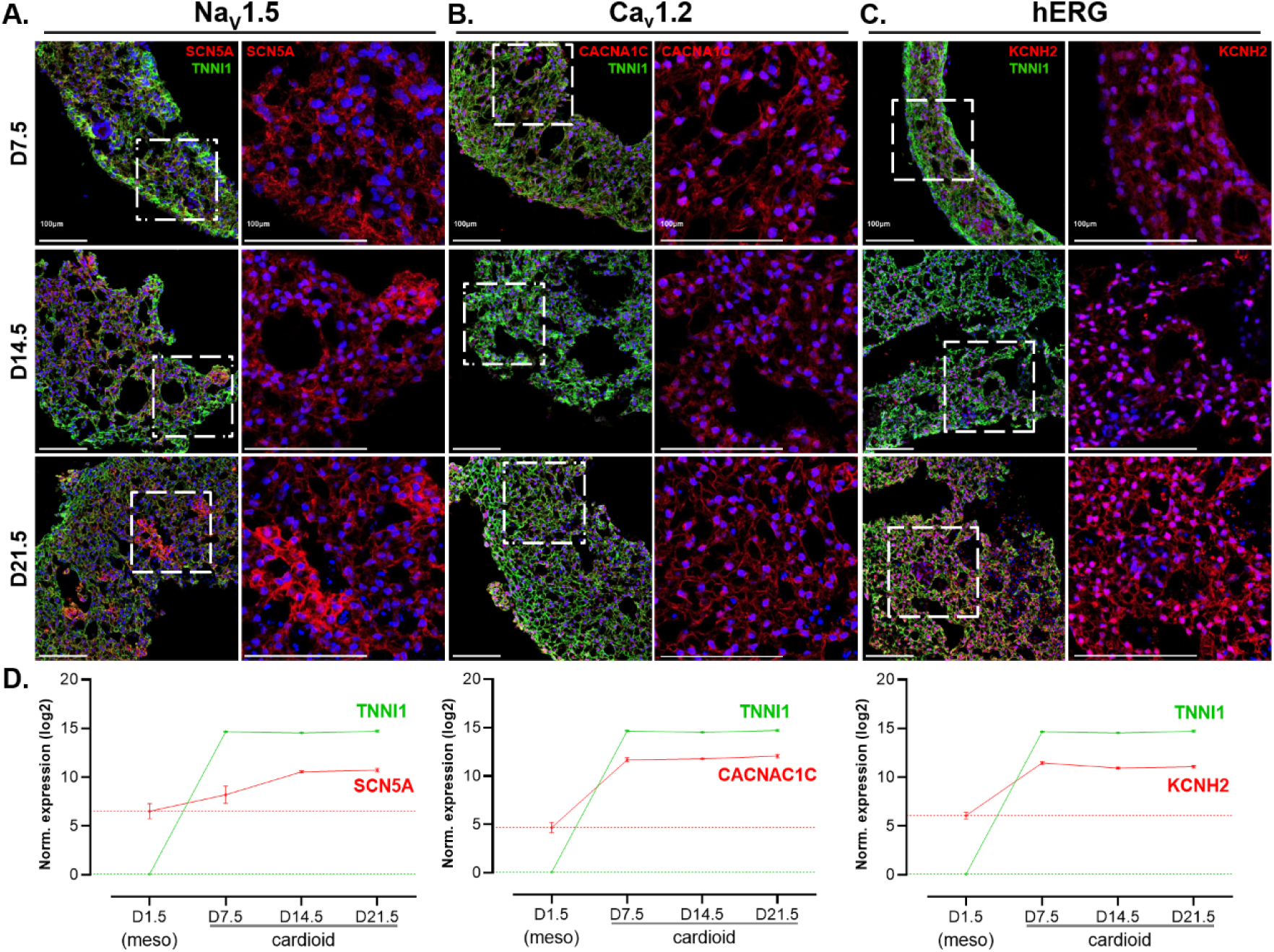
Expression dynamics of key depolarizing and repolarizing cardiac ion channels in maturated cardioids supports the electrophysiological characterization. **A-C.** Representative immunofluorescence results (n=3/3) demonstrating increased protein expression levels of Na_v_1.5 (A), hERG (B) and Ca_v_1.2 (C) in cardioids at day 7.5, 14.5 and 21.5. Panels on the left represent magnified region within the dashed outline. Scale bars, 100μm. **D.** Log2-normalized RNA expression profiles (n=3 biological replicates) of *SCN5A*, *CACNA1C*, and *KCNH2* ion channels, plotted relative to *TNNI1* across key developmental time points (D1.5, D7.5, D14.5, and D21.5), demonstrating transcriptomic signatures consistent with observed functional protein expression.

Na_v_1.5 immunoreactivity increased progressively during cardioid maturation (**Figure 6A**), consistent with the observed increase in I_Na_ current density (**Figure 5A**). At D14.5, and more prominently at D21.5, regions containing CM clusters with high Nav1.5 immunoreactivity became apparent (**Figure 6A**). These Nav1.5-high regions were larger and more spatially dispersed within the whole cardioids at D21.5. Consistent with the protein-level observations, *SCN5A* transcript abundance increased during maturation, in particular from D7.5 to D14.5 (log2FC: 1.802; P=3.56E-04) (**Figure 6D**).

Our RNA-seq data also revealed that *KCND2* and *KCND3* transcript levels (e.g., the genes encoding K_v_4.2 and K_v_4.3 channels that contribute to I_to_) were relatively low but increased progressively during cardioid maturation (**Suppl. Fig. 5D**). Nevertheless, I_to_ was detected throughout cardioid maturation (**Suppl. Fig. 5A**), with no significant changes in current density across the examined time points.

hERG immunoreactivity and *KCNH2* transcript abundance remained relatively stable throughout maturation (**Figure 6B & D**), consistent with stable I_Kr_ current density (**Figure 5C**). Similarly, Ca_v_1.2 immunoreactivity and *CACNA1C* transcript abundance were comparable across D7.5, D14.5, and D21.5, albeit I_CaL_ current densities slightly increased with maturation (**Figure 5B**).

*KCNQ1* transcripts were readily detected throughout maturation (**Suppl. Fig. 5B**). However, mRNA expression of the auxiliary β-subunit *KCNE1* remained comparatively low (**Suppl. Fig. 5D**), which may contribute to the limited detection of I_Ks_ in isolated cardioid-derived cardiomyocytes despite its proven functional relevance to shape AP duration in intact cardioids (**Suppl. Fig. 4B & 4D**).

Interestingly, *KCNJ2* mRNA expression increased during maturation from week 1 to 2 (log2FC: 5.551; P=3.16E-12) as well as from week 2 to week 3 (log2FC: 1.866; P=5.75E-10) – in line with the progressively more negative RMP detected in intact cardioids, but remained lower than *SCN5A*, *CACNA1C*, and *KCNH2* (**Suppl. Figure 5D**). Remarkably, among the ion channels examined, only *KCNJ2* and *KCNJ3* were upregulated in both the D7.5-to-D14.5 and D14.5-to-D21.5 comparisons (**Suppl. Fig. 6, Suppl. Table 5**). In support, cardiomyocytes in slices of D21.5 cardioids showed stronger Kir2.1 signal than at D7.5, with a subcellular distribution like that observed in adult rabbit ventricular sections, but overall lower signal than in the ventricular tissue (**Suppl. Fig. 7**).

Together, these findings demonstrate progressive electrophysiological maturation of human cardioids during the first three weeks of development and reveal concordance between tissue-level electrophysiology and calcium handling, ion current characteristics and molecular profiling.

## Discussion

In the present study, we combined sharp microelectrode recordings, calcium imaging, whole-cell patch clamp recordings, pharmacological interventions, immunofluorescence analyses, and transcriptomic profiling to provide the first integrated, comprehensive electrophysiological characterization of human cardioids over three weeks of maturation. Our results demonstrate that cardioids undergo progressive functional maturation, characterized by progressively more negative resting membrane potentials, improved action potential characteristics and progressively increased I_Na_ and I_CaL_ current densities, as well as persistent functional repolarizing I_Kr_ currents. However, several electrophysiological properties remain immature, most notably the persistent absence of detectable I_K1_ in isolated cardioid-derived cardiomyocytes despite mature RMP on the intact cardioid level.

### Progressive electrophysiological maturation in cardioids

The principal finding of this study is that 3D cardioids undergo progressive electrophysiological maturation during the first three weeks of culture. RMP became more negative, while APA and dV/dt_max_ increased with maturation (**Figure 2**), approaching values reported for adult human ventricular myocardium, where RMP is approximately −85 mV, APA is >100 mV (Jost et al., 2013a; Mohammed et al., 2026) and dV/dt_max_ is approximately 200-250 mV/ms (Lemoine et al., 2017). These values were obtained using conventional sharp microelectrode recordings, providing a relevant tissue-level comparison for our cardioid recordings.

These changes in AP characteristics were accompanied by progressively faster calcium transient kinetics (**Figure 4**), indicating that electrical maturation was associated with improved calcium handling (Sun et al., 2024). Similar progressive maturation of hiPSC-CMs has been reported during prolonged culture, particularly when cells are exposed to structural or multicellular cues such as co-culture, micropatterning, or 3D environments (Lundy et al., 2013; Lewandowski et al., 2018; Seibertz et al., 2023). Several processes may contribute to this maturation, including improved sarcomere organization, enhanced cell-cell coupling, metabolic maturation, increased ion channel expression, and more efficient excitation-contraction coupling (Bedada et al., 2016; Dai et al., 2017; Huang et al., 2020; Lundy et al., 2013).

Cardioids provide a 3D multicellular environment that more closely resembles native cardiac tissue than conventional 2D monolayers. The close spatial organization of cardiomyocytes and non-myocyte populations may facilitate electrical coupling and synchronized excitation (Xiao et al., 2026), potentially contributing to the less variable action potential recordings observed at later maturation stages. Although cardioids remain developmentally immature, these findings demonstrate that substantial electrophysiological maturation occurs within only three weeks of culture.

### Sodium channel maturation improves cardiac excitability

The improvement in AP characteristics closely paralleled maturation of the cardiac sodium current. Peak I_Na_ current density increased markedly between weeks 1 and 3 (**Figure 5A**), accompanied by increased Na_v_1.5 protein expression (**Figure 6A**) and progressively elevated *SCN5A* transcript levels (**Figure 6D**), demonstrating coordinated maturation at the transcript, protein, and functional levels. In addition to the increase in peak I_Na_ density, maturation was accompanied by a progressive shift in the distribution of sodium current phenotypes. The proportion of cardiomyocytes lacking measurable I_Na_ decreased throughout maturation, whereas an increasing proportion of cells displayed very large I_Na_ currents that could not be adequately voltage-clamped (**Suppl. Fig. 4A**).

The increase in I_Na_ is expected to enhance membrane excitability and contribute to a faster AP upstroke and larger AP amplitude. Previous studies have linked increased I_Na_ current density to increased AP upstroke velocity in hiPSC-CMs and 3D cardiac tissues (Lemoine et al., 2017; Seibertz et al., 2023). Consistent with this, the increase in I_Na_ in our cardioids was accompanied by a progressive increase in APA and upstroke velocity (**Figure 2, Suppl. Table 1**).

### Functional repolarization develops early despite incomplete maturation

In contrast to sodium currents, major repolarizing potassium currents were already functionally established in cardioids at early stages. Pharmacological inhibition of I_Kr_ with dofetilide consistently prolonged APD_80_ in cardioids across the examined weeks (**Figure 3A**), consistent with stable hERG protein expression (**Figure 6C**), *KCNH2* transcript (**Figure 6D**), and I_Kr_ current density (**Figure 5C**). These findings highlight the suitability of cardioids for studying *KCNH2*-related repolarization disorders and for evaluating hERG-mediated pro-arrhythmic drug effects (Yaqinuddin et al., 2025).

Similarly, I_Ks_ contributed to tissue-level repolarization, as demonstrated by significant APD_80_ prolongation following HMR-1556 application alone or after I_Kr_-inhibition induced reduced repolarization reserve (**Figure 3B, Suppl. Fig 3**). This is consistent with previous studies (Jost et al., 2007; Jost et al., 2013) showing that I_Ks_ serves as important component of repolarization reserve – particularly under conditions of impaired repolarization. In contrast, isolated cardioid-derived cardiomyocytes exhibited only small steady-state I_Ks_ currents, lacked clearly detectable tail currents, and did not consistently express I_Ks_ in all cells (**Suppl. Fig. 5B**). Importantly, although *KCNQ1* and *KCNE1* transcripts were detected throughout weeks 1–3, *KCNE1* expression remained substantially lower than *KCNQ1* in the cardioids (**Suppl. Fig. 5D**). Heterologous expression studies have shown that *KCNQ1* with insufficient *KCNE1* results in rapidly activating currents that also undergo a rapid inactivation, thus lacking the characteristic slowly activating I_Ks_ with prominent tail currents observed upon *KCNQ1*-*KCNE1* co-expression (reviewed in depth in Wu et al., 2020). Thus, the low *KCNE1/KCNQ1* expression ratio in cardioids may explain why isolated cardioid-derived cardiomyocytes exhibited only small steady-state outward currents without clearly detectable I_Ks_ tail currents. Similar difficulties in reliably detecting I_Ks_ have been reported previously in hiPSC-derived cardiomyocytes (Ma et al., 2015; Li et al., 2025).

### Tissue-level electrophysiology reveals properties not readily detected in isolated cardioid-derived cardiomyocytes

A key finding of this study is the difference between electrophysiological measurements obtained from intact cardioids and isolated cardioid-derived cardiomyocytes, highlighting the importance of evaluating electrophysiological properties and pharmacological responses within an intact 3D tissue architecture. Pharmacological experiments in intact cardioids demonstrated clear effects of HMR-1556 and BaCl₂, indicating a functional contribution of I_Ks_ to repolarization and of barium-sensitive currents to RMP regulation and repolarization. In contrast, whole-cell patch clamp recordings in isolated cardioid-derived cardiomyocytes detected only minimal I_Ks_ and no I_K1_ current.

For I_Ks_, 58% of isolated patched cardioid-derived cardiomyocytes displayed a measurable steady-state I_Ks_, whereas the remaining cells showed no detectable I_Ks_ under our experimental conditions. This is consistent with the heterogeneous expression of I_Ks_ previously reported in hiPSC-CMs (Ma et al., 2011; Moretti et al., 2010; Hoekstra et al., 2012). In intact cardioids, electrotonic coupling between cardiomyocytes may allow such heterogeneous cellular currents to contribute to the integrated tissue-level response, as illustrated by the robust APD prolongation induced by HMR-1556 (Figure 3B).

A similar divergence was observed for the barium-sensitive response. BaCl₂ depolarized the RMP, reduced APA, and prolonged repolarization in intact cardioids, whereas a classical I_K1_ current was not detected in isolated cardioid-derived cardiomyocytes. The distinct nature of the two recording approaches may contribute to this difference: sharp microelectrode recordings measure the integrated electrical behaviour of an electrotonically coupled syncytium, whereas patch-clamp recordings assess individual cardiomyocytes. Consequently, small membrane conductance distributed across interconnected cardiomyocytes may substantially influence tissue electrophysiology while remaining undetectable in isolated cells (Horváth et al., 2018). Consistent with this, computational and experimental studies suggest that only modest I_K1_ is required to stabilize the RMP in multicellular preparations (Reilly & Eckhardt, 2021; Dhamoon & Jalife, 2005). In addition, enzymatic isolation may alter cellular electrophysiology by disrupting membrane integrity, damaging ion channels, and abolishing native cell-cell interactions (Koc et al., 2021). Furthermore, cardioids contain heterogeneous cardiomyocyte populations at different maturation states (Hofbauer et al., 2021), so that small subpopulations expressing functional ion channels may contribute to tissue-level electrophysiology, while being underrepresented in single-cell recordings. In line with this, we observed progressively increased *KCNJ2* transcript during maturation but detected only limited Kir2.1 protein expression in the cardioids, which may have contributed to lack of clearly detectable I_K1_ at the cellular level (**Suppl. Fig 7**).

The BaCl₂ response should however be interpreted with caution. Despite being is widely used to inhibit I_K1_, BaCl₂ is not completely selective and its effects depend on concentration and membrane potential. In cardiac preparations, Ba²⁺ can also inhibit other potassium currents, including I_to_ (Dhamoon & Jalife, 2005), although the sensitivity of I_K1_ is substantially higher at negative membrane potentials (Nerbonne, 2016). Since I_to_ contributes primarily to early action potential repolarization, it is unlikely to account for the changes in APD_80_ and RMP observed following BaCl₂ application (**Figure 3C**). Other constituent background potassium conductances may contribute to the BaCl_2_ response observed in intact cardioids (Jeevaratnam et al., 2018), most notably two-pore domain background potassium leak channels (K_2P_, e.g., TASK channels), as well as low-level basal activation of calcium-activated potassium conductances (Zhou et al., 2012; Rothberg, 2012; Warren et al., 2003b). Therefore, while the observed changes in RMP and repolarization are consistent with inhibition of an I_K1_-like conductance, our findings demonstrate the presence of functional barium-sensitive conductances contributing to tissue electrophysiology without establishing the exclusive molecular identity of the underlying current.

Together, these findings highlight the complementary information provided by tissue- and single-cell electrophysiological recordings. Preserving multicellular architecture and electrical coupling in cardioids may allow relatively small or heterogeneous conductance to exert measurable effects at the tissue level despite being difficult to detect consistently in isolated cardiomyocytes.

### Calcium handling matures together with membrane electrophysiology

Electrophysiological maturation was accompanied by enhanced I_CaL_ and improved calcium handling, reflected by faster calcium transient upstrokes and a shorter time to peak in calcium transients in line with the progressively enhanced I_CaL_. Moreover, action potential and calcium transient duration shortened in parallel during maturation, consistent with improved excitation–contraction coupling. Although calcium cycling proteins were not directly quantified, these functional changes are consistent with progressive maturation of sarcoplasmic reticulum calcium handling, including improved SERCA activity, ryanodine receptor function (RYR2), and sodium-calcium exchanger regulation (NCX), as previously described during hiPSC-CM maturation (Hwang et al., 2015). The parallel improvement of membrane electrophysiology and calcium handling further supports coordinated functional maturation of cardioids rather than isolated changes in individual ion channels.

### Implications for disease modelling

An important objective in developing advanced human cardiac models is their application in cardiac channelopathy modelling and safety pharmacology. The present study demonstrates that, following maturation, cardioids exhibit adult-like resting membrane and action potentials, mature I_Na_, functional calcium handling, and stable I_Kr_, making them well suited for studying disorders of depolarization and repolarization, including *KCNH2*-mediated long and short QT syndromes and sodium channelopathies. Importantly, electrophysiological recordings in intact 3D cardioids preserve 3D multicellular electrical interactions absent in isolated cells and therefore more closely reflect native cardiac physiology and pathophysiology (Jæger et al., 2025).

### Study limitations

Several limitations should be acknowledged. First, electrophysiological characterization was limited to the first three weeks of maturation, and longer culture periods may reveal further maturation. Second, although patch clamp recordings were performed 16–48 h after cardioid dissociation - earlier than in most previous studies (Hofbauer et al, 2021; Schmidt et al, 2022) - to minimize dedifferentiation and electrophysiological alterations associated with prolonged 2D culture (Koc et al., 2021), dissociation-induced membrane damage may still have influenced ion channel function. Finally, cardioids contain heterogeneous cardiomyocyte populations, and the relative contributions of ventricular-, atrial-, and nodal-like cells were not examined. Importantly, however, all tissue-level and single-cell recordings were performed on cardioids from the same differentiation batches, ensuring biological comparability across experimental modalities.

## Conclusion

In conclusion, this study provides the first integrated, comprehensive electrophysiological characterization of human cardioids and their maturation by integrating tissue-level electrophysiology with single-cell ion channel recordings, calcium imaging, molecular protein expression and transcriptomic profiling, and pharmacological analyses. Human cardioids exhibit progressive electrophysiological maturation, characterized by mature AP characteristics, including a progressively more negative RMP, enhanced sodium and calcium channel function, accelerated calcium handling, and stable I_Kr_-mediated repolarization. At the same time, the persistent absence of detectable classical cellular-level I_K1_ highlights an important remaining limitation shared with other hiPSC-derived cardiac models. Overall, our findings establish cardioids as a robust and scalable three-dimensional platform for investigating human cardiac electrophysiology, disease mechanisms, and drug responses while providing a framework for future strategies aimed at achieving fully mature human cardiac tissue.

## Acknowledgements

We are grateful to Sasha Mendjan and Stefan M. Jahnel for training and guidance in establishing LV cardioids. We thank Dr. Jean-Sébastien Rougier for valuable support and discussions regarding electrophysiological experiments and for kindly providing sodium-antibodies. We also thank Dr. Nina Ullrich for valuable advice and discussions regarding hiPSC culture, as well as for kindly providing calcium-related antibodies. We also thank Julien Louradour for his valuable comments and suggestions on the manuscript. This project was funded by a Bern Center of Precision Medicine (BCPM) Lighthouse grant to K.E.O. and M.O. M.O. and I.B. were additionally supported by NRP79 grant #206520 from the Swiss National Science Foundation.

---

**Suppl. Fig. 1.**
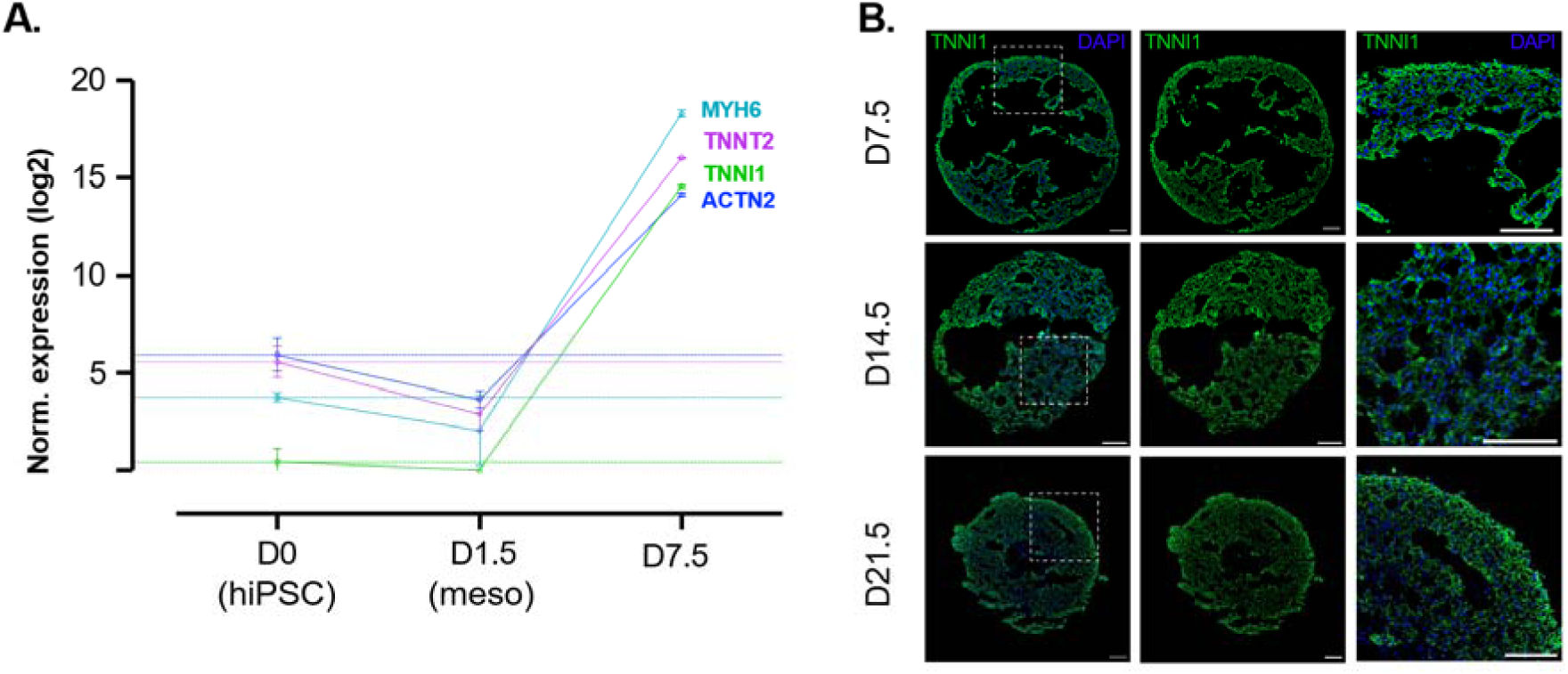
Molecular characterization of LV-type cardioids. B. Line plot showing RNA-seq expression profiles of the cardiomyocyte marker genes *MYH6*, *TNNT2*, *TNNI1* and *ACTN2* in hiPSCs (D0), mesodermal progenitors (D1.5), and cardioids at D7.5. Dashed lines indicate baseline levels. **B**. Anti-GFP immunofluorescence in cardioids generated from the TNNI1-GFP WTC11 hiPSC line at D7.5, D14.5 and D21.5. GFP signal identifies *TNNI1*-positive cardiomyocytes, which were previously shown to comprise approximately 90% of the cells in LV cardioids (Hofbauer et al., 2021). Dashed boxes in the left panels delineate the regions shown at higher magnification in the corresponding right panels. Scale bars, 100μm.

**Suppl. Fig. 2.**
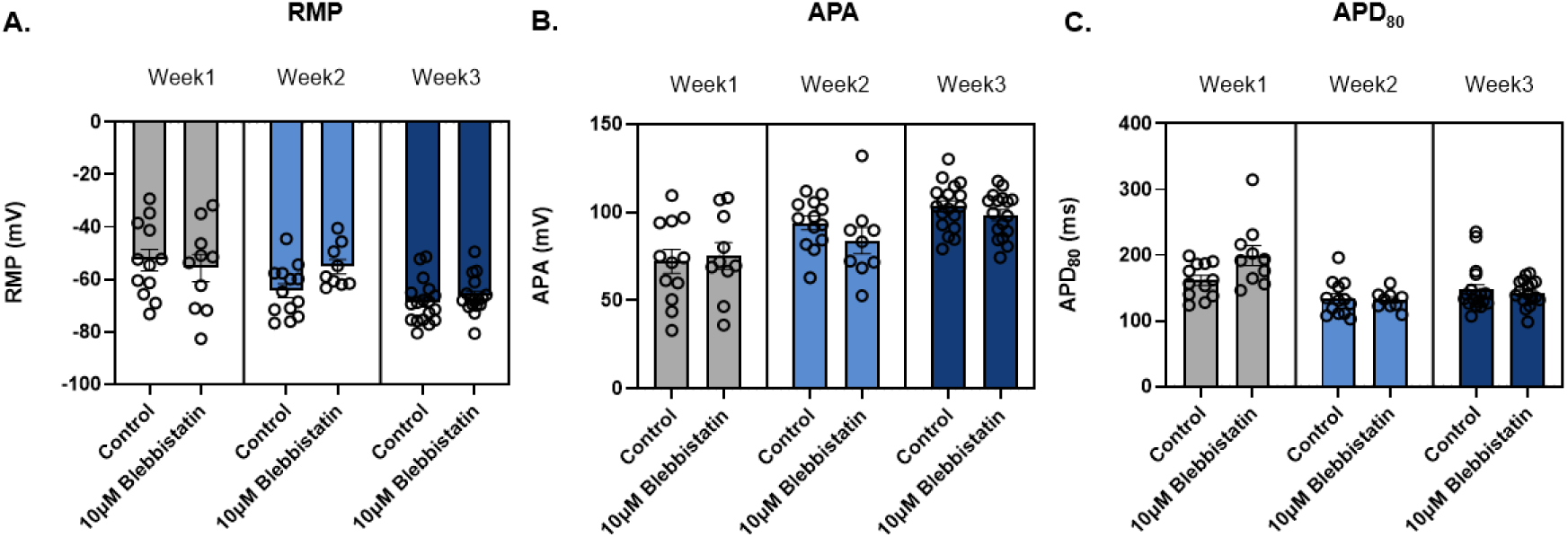
Comparison of action potential characteristics in cardioids recorded under control conditions and following incubation with 10 μM blebbistatin. A. Resting membrane potential (RMP). B. Action potential amplitude (APA). C. Action potential duration at 80% repolarization (APD_80_) at weeks 1, 2, and 3. Statistical significance was defined as *p*<0.05.

**Suppl. Fig. 3.**
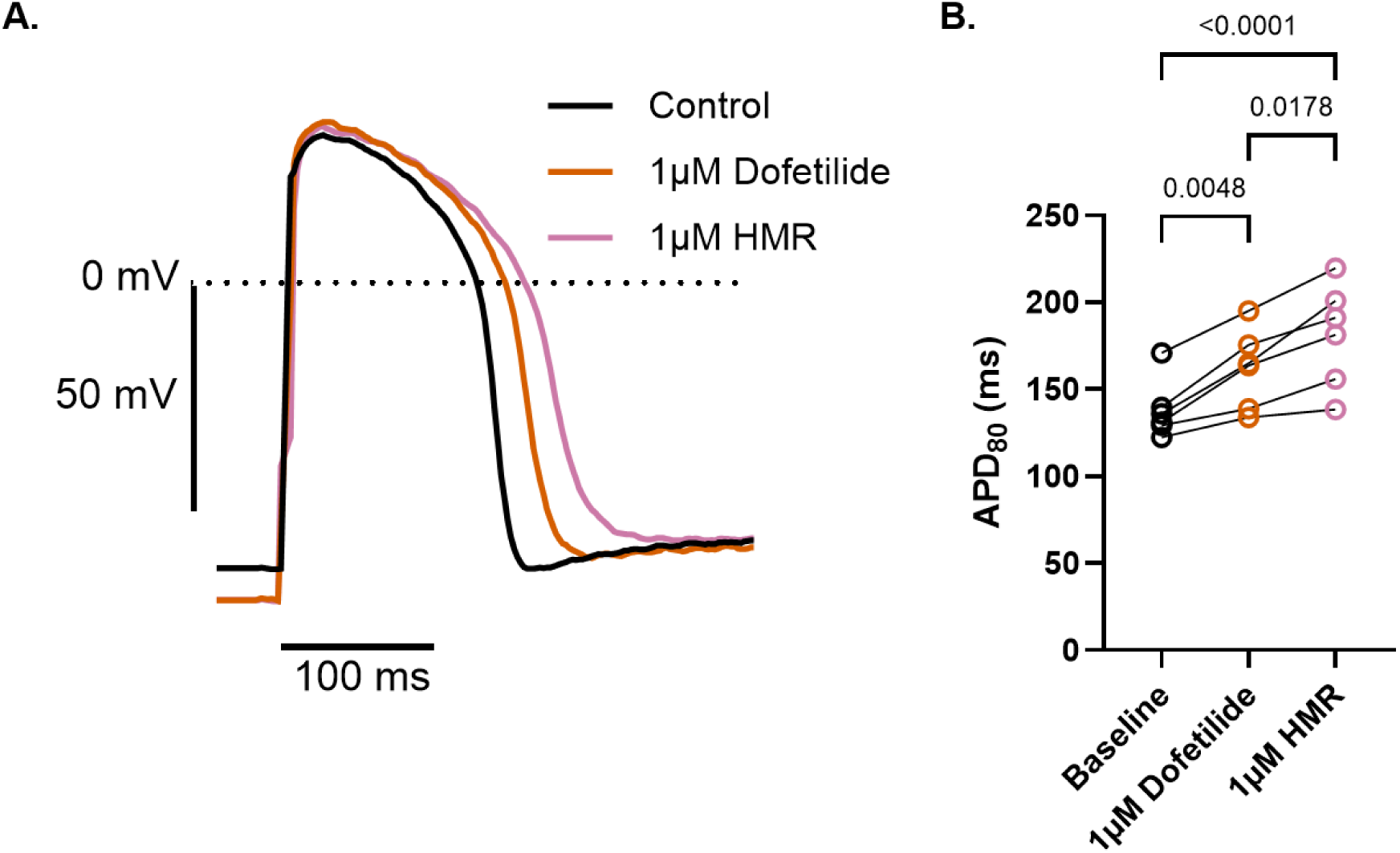
Sequential pharmacological inhibition of I_Kr_ and I_Ks_ in intact cardioids. Dofetilide (1 μM) was first applied to cardioids from week 2 (n=2) and week 3 (n=3), resulting in a significant prolongation of APD_80_. HMR-1556 (1 μM) was subsequently applied in the continued presence of dofetilide, resulting in a further prolongation of APD_80_ compared with dofetilide alone (*p*<0.05), indicating an additional contribution of I_Ks_ to repolarization. Data are presented as mean ± SEM. Statistical significance was assessed using a repeated-measures one-way ANOVA. *p*<0.05.

**Suppl. Fig. 4.**
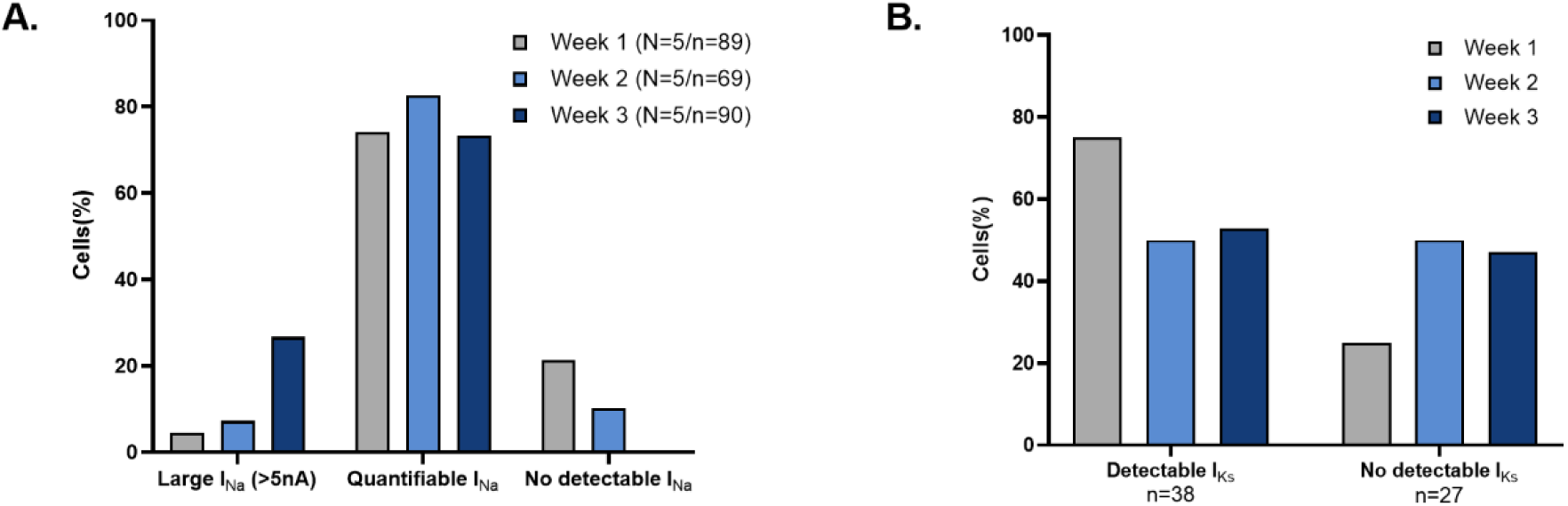
Additional analyses supporting electrophysiological maturation of cardioids. **A.** Percentage of cardioid-derived cardiomyocytes exhibiting excessively large sodium currents (>5 nA), quantifiable sodium currents, or no detectable sodium currents at weeks 1, 2, and 3. **B.** Percentage of patched cardioid derived CMs exhibiting detectable I_Ks_ (58.5%) or no detectable I_Ks_ (41.5%) throughout the maturation weeks.

**Suppl. Fig. 5.**
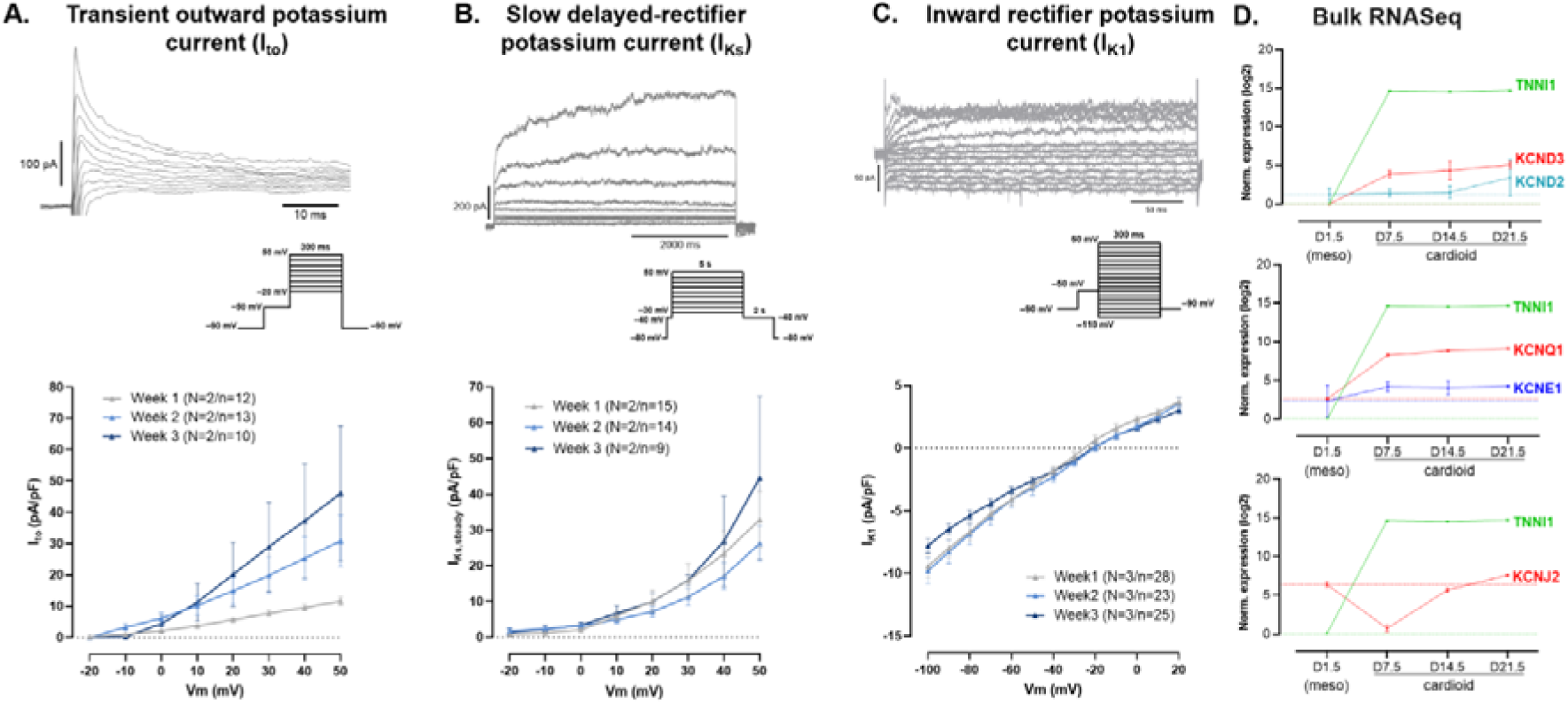
Changes in cardiac ion currents during cardioid maturation. **A.** Representative I_to_ current traces (top) and current-voltage relationships (bottom) recorded from cardioid-derived cardiomyocytes at weeks 1, 2, and 3. **B.** Representative I_Ks_ current traces (top) and steady-state current-voltage relationships (bottom). **C.** Representative I_K1_ current traces (top) and current-voltage relationships (bottom). **D.** Bulk RNA-sequencing analysis showing log2-normalized expression of *KCND2/KCND3* (I_to_), *KCNQ1/KCNE1* (I_Ks_), and *KCNJ2* (I_K1_), plotted relative to the cardiac maturation marker TNNI1 across key time points (D1.5, D7.5, D14.5 and D21.5).

**Suppl. Fig. 6.**
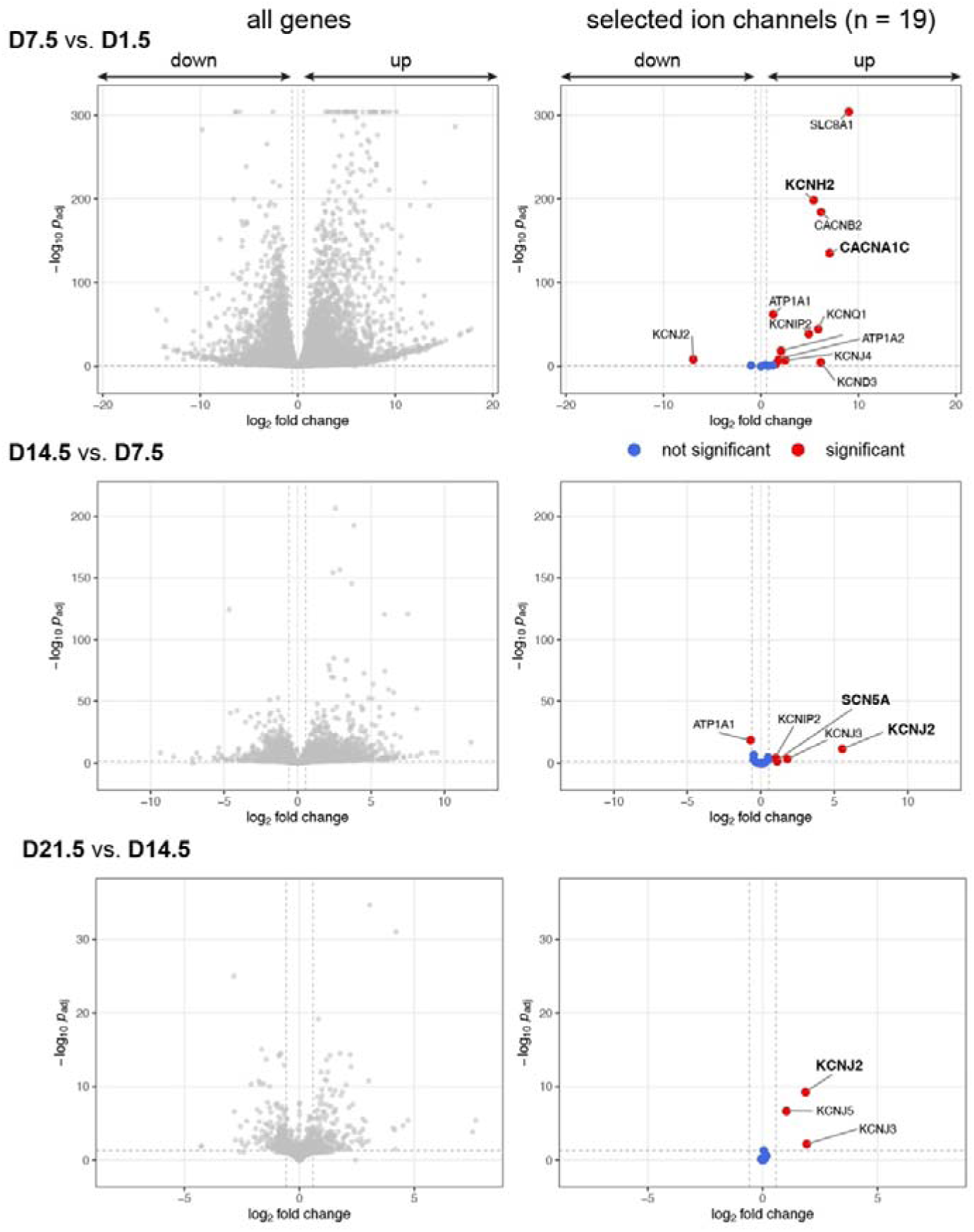
Differentially expressed ion channels during maturation of LV-type cardioids. DESeq2 analysis of ion channel genes differentially expressed between the mesodermal progenitor and mature cardioid stages (D1.5 vs. D7.5), as well as across early and late cardioid maturation (D7.5 vs. D14.5 and D14.5 vs. D21.5). Left panels show all differentially expressed genes and right panels highlight ion channels from a curated list relevant to LQTS (Suppl. Table 5).

**Suppl. Fig. 7.**
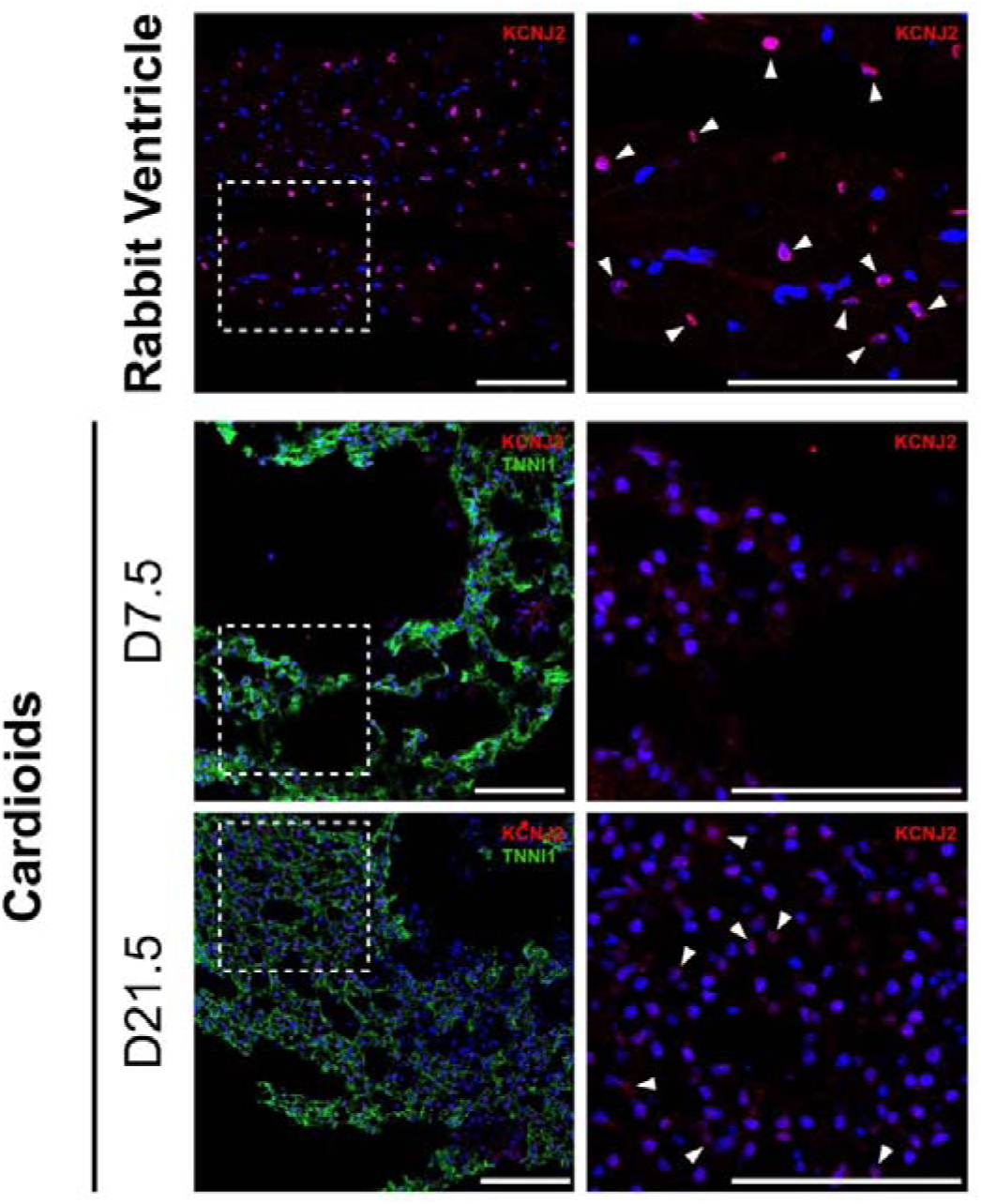
Kir2.1 antibody staining in rabbit cardiac ventricle and LV-type cardioids. Anti-GFP and anti-Kir2.1 immunofluorescence staining of a rabbit ventricular heart section (top panels) and cardioids generated from the TNNI1-GFP WTC11 hiPSC line at days 7.5 and 21.5, (bottom panels). GFP (green) marks TNNI1-positive cardiomyocytes, and Kir2.1 staining is shown in red. Dashed boxes in the left panels indicate regions shown at higher magnification in the corresponding right panels. Arrowheads point to localization of Kir2.1 signal. Scale bars, 100 μm.

## Supplemental data

**Suppl. Table 1.**
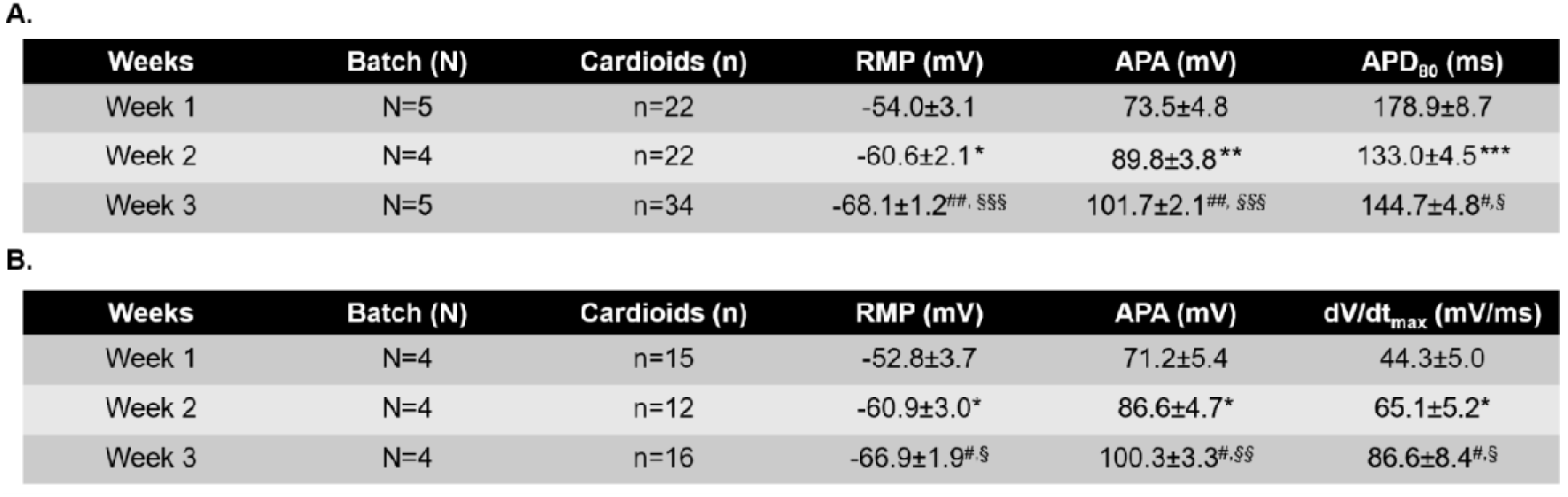
Baseline action potential characteristics of cardioids at different maturation stages A. Baseline action potential parameters including resting membrane potential (RMP), action potential amplitude (APA), and action potential duration at 80% repolarization () recorded across weeks 1, 2, and 3 using conventional sharp microelectrodes. **B.** Maximum upstroke velocity (dV/dt_max_) measured in a subset of action potential recordings free from pacing artifact interference, alongside corresponding RMP and APA values. Results are presented as mean ± SEM. Statistical comparisons were performed using one-way ANOVA or the Kruskal-Wallis test, as appropriate. *\*p*<0.5, *\*\*p*<0.01, *\*\*\*p*<0.0001 for week 1 vs. week 2; *^#^p*<0.5, *^##^p*<0.05, *^###^p*<0.0001 for week 2 vs. week 3; *^§^p*<0.01, *^§§§^p*<0.0001 for week 1 vs. week 3.

**Suppl. Table 2.** Characteristics of calcium transients in intact cardioids at different maturation stages. Time to peak, Upstroke velocity (dcdt, ΔF/F0*ms-1), and calcium transient duration at 80% recovery (CaD_80_) were measured in intact cardioids at weeks 1, 2, and 3. Results are presented as mean ± SEM. N indicates the number of independent differentiation batches and n the number of cardioids. Statistical comparisons were performed using one-way ANOVA or the Kruskal-Wallis test, as appropriate. *\*\*\*p*<0.0001 for week 1 vs. week 2; *^##^p*<0.01 for week 2 vs. week 3; *^§^p*<0.05, *^§§^p*<0.01, *^§§§^p*<0.0001 for week 1 vs. week 3.

| Weeks | Batch (N) | Cardioids (n) | Time to peak (ms) | dcdt (ΔF/F<sub>0</sub>*ms<sup>-1</sup>) | CaD<sub>80</sub> (ms) |
| --- | --- | --- | --- | --- | --- |
| Week1 | N=3 | n=10 | 315.8±12.8 | 3.0±0.2 | 790.0±13.4 |
| Week2 | N=3 | n=10 | 199.4±11.9*** | 7.4±0.4*** | 655±15.8*** |
| Week3 | N=3 | n=10 | 215.4±10.5<sup>SSS</sup> | 4.9±0.6<sup>##</sup>, <sup>SS</sup> | 720.9±14.5<sup>S</sup> |

**Suppl. Table 3. RNA-seq sample metadata and quality-control metrics (provided as excel file).** Samples are identified by differentiation day (D) and biological replicate (timepoint_replicate); original stage labels, sequencing batches and nf-core/rnaseq sample IDs are provided for traceability. The table summarizes raw-read quality, trimming, STAR alignment, post-alignment metrics and Salmon quantification. Read counts refer to reads for single-end libraries and read pairs for paired-end libraries; percentages are calculated using the indicated raw, post-trimming, alignment-input or biotype-count denominator. Mate-level trimming and GC metrics are averaged for paired-end libraries. Each sequencing batch was processed in a separate nf-core/rnaseq run.

**Supplementary Table 4. RNA-seq analysis (provided as excel file).** Matrices providing gene-level estimated raw counts, DESeq2-normalized counts without batch correction, log2-transformed normalized expression values after batch correction (bc), and variance-stabilized counts (VST) before and after removal of the sequencing batch effect.

**Supplementary Table 5. Differential expression analysis of selected ion channels during cardioid differentiation (provided as excel file).** Sheets 1-2: Summary of transcriptome-wide differential expression across pairwise comparisons of D0 (hiPSCs), D1.5 (mesodermal progenitors), and D7.5, D14, and D21.5 cardioids. Sheets 3-4: Log₂ fold changes (log₂FC) and adjusted P values (padj) for 19 selected ion channels. Sheets 5–10: Differentially expressed genes for the D7.5 versus D1.5, D14.5 versus D7.5, and D21.5 versus D14.5 comparisons.

